# Prostaglandin E_1_ as therapeutic molecule for Nephronophthisis and related ciliopathies

**DOI:** 10.1101/2022.01.21.477191

**Authors:** Hugo Garcia, Alice Serafin, Flora Silbermann, Esther Poree, Clémentine Mahaut, Amandine Viau, Katy Billot, Éléonore Birgy, Meriem Garfa-Traore, Stéphanie Roy, Salomé Cecarelli, Manon Mehraz, Pamela C. Rodriguez, Bérangère Deleglise, Laetitia Furio, Fabienne Jabot-Hanin, Nicolas Cagnard, Elaine Del Nery, Marc Fila, Soraya Sin-Monnot, Corinne Antignac, Stanislas Lyonnet, Pauline Krug, Rémi Salomon, Jean-Philippe Annereau, Alexandre Benmerah, Marion Delous, Luis Briseño-Roa, Sophie Saunier

**Author notes:** the three first authors participate equally to the work (Hugo Garcia, Alice Serafin, Flora Silbermann). the co-last authors contribute equally to the work (Alexandre Benmerah, Marion Delous, Luis Briseño-Roa, Sophie Saunier).

## Abstract

Nephronophthisis (NPH) is an autosomal recessive tubulointerstitial nephropathy belonging to the ciliopathy disorders and known as the most common cause of hereditary end-stage renal disease in children. Yet, no curative treatment is available. The major gene, *NPHP1*, encodes a protein playing key functions at the primary cilium and cellular junctions. Using an *in cellulo* medium-throughput drug-screen, we identified 51 FDA-approved compounds and selected 11 for their physicochemical properties, including prostaglandin E_1_ (PGE1). PGE1 was further validated to rescue ciliogenesis in immortalized patient *NPHP1^-/-^* urine-derived renal tubular cells and corroborated by the effects of its analog PGE2. The two molecules reduced pronephric cyst occurrence *in vivo* in *nphp4* zebrafish model, and PGE1 treatment in *Nphp1^-/-^* mice led to a significant reduction of renal tubular dilatations, partially restoring cilia length within tubules. Finally, comparative transcriptomics allowed identification of key molecules downstream PGE1. Altogether, our drug-screen strategy led to the identification of PGE1 as the first potential therapeutic molecule for NPH-associated ciliopathies.

**Significant statement:** Juvenile nephronophthisis (NPH) is a renal ciliopathy due to a dysfunction of primary cilia and a common genetic cause of end-stage renal disease in children and young adults. No curative treatment is available. This paper describes the identification of Prostaglandin E1 (PGE1) as the first potential therapeutic molecule for NPH-associated ciliopathies. We demonstrated that PGE1 rescues defective ciliogenesis and ciliary composition in *NPHP1^-/-^* patient urine-derived renal tubular cells. Furthermore, PGE1 improves ciliary and kidney phenotypes in our NPH zebrafish and *Nphp1^-/-^* mouse models. Finally, *in vitro* experiments as well as transcriptomic analyses pointed out several pathways downstream PGE1 as cAMP, cell-cell/cell-matrix adhesion or actin cytoskeleton. Altogether, our findings provide a new alternative for treatment of NPH.

## Introduction

Renal ciliopathies are a group of hereditary diseases caused by mutations in genes encoding for proteins playing a role at the primary cilium, a microtubule-based cellular antenna found on almost all vertebrate cells which controls key signaling pathways during development and tissue homeostasis (1). In the kidney, as well as in other organs, the primary cilium also acts as an essential mechanosensor through which luminal flow controls tubule diameter and renal epithelium maintenance, explaining the presence of tubular cysts in most renal ciliopathies. Renal ciliopathy spectrum includes autosomal dominant polycystic kidney disease (ADPKD), autosomal recessive polycystic kidney disease and cystic dysplasia, which are all characterized by enlarged kidneys with numerous cysts (2). It also encompasses nephronophthisis (NPH) that, on the other hand, is generally characterized by small kidneys and progressive tubulointerstitial fibrosis, associated with corticomedullary cysts, leading to end-stage renal disease (ESRD) at a median age of 12 years-old. NPH is the first monogenic cause of pediatric ESRD, representing up to 10% of incident cases in developed countries (3, 4). Recent publications have demonstrated that NPH may also occur in adulthood, leading to late onset ESRD (up to 61 years-old) (5). Extrarenal involvements may include retinal degeneration (Senior-Løken syndrome), cerebellar ataxia, skeletal anomalies, and/or liver fibrosis (6).

To date, more than 23 genes have been linked with NPH, the most frequent being *NPHP1* with nearly 25% of affected individuals harboring a homozygous deletion of this gene. *NPHP* genes encode proteins that form different complexes along the cilium (7). Their mutations are not associated with a complete loss of cilia; they rather lead to subtle phenotypes such as decreased proportion of ciliated cells increased cilium length and/or ciliary membrane composition defects which result in defective cilia-dependent signaling (8–10). Besides cilium dysfunction, several NPH-associated gene products, including NPHP1 and NPHP4, have also been involved in the control of cell migration, adhesion and apico-basal cell epithelialization, through the regulation of cytoskeletal organization (8, 11–14).

To date, the only available treatments for NPH patients are dialysis and renal transplantation. As other chronic tubulointerstitial nephropathies characterized by the absence of hypertension or overt albuminuria (except in late stages of the disease), non-specific nephroprotective treatments by renin-angiotensin system blockade did not delay progression to ESRD (15). Similarly, as enlarged cystic kidneys are uncommon in juvenile NPHP1-associated NPH, molecules developed to limit cyst growth in the context of ADPKD such as the selective angiotensin-vasopressin type 2 (AVPR2) antagonist Tolvaptan, the CDK inhibitor Roscovitine or the somatostatin-analog Octreotide are unlikely to be relevant (16, 17). Nevertheless, several other molecules were recently pointed out in *in vitro* and *in vivo* NPH models including the Hedgehog agonist Purmorphamine, β-catenin inhibitors, ROCK inhibitors or the flavonoid Eupatilin (4). On the other hand, even though gene therapy has been successfully implemented in early human retinal ciliopathies thanks to local administration of adeno-associated viruses (18), efficient gene therapy in NPH would rather represent a clinical challenge given the lack of effective administration modes for gene delivery in the kidney.

Phenotypic medium-throughput screening strategies have already led to the identification of a potential therapeutic molecule in the context of CEP290/NPHP6 (19). Here, we performed *in vitro* screens of small chemical compounds for their ability to lessen phenotypes associated with the loss of *NPHP1.* These screens led to the identification of prostaglandin E_1_ (PGE1)-related compounds as a new class of molecules that restore ciliary defects *in vitro* and *in vivo* in both zebrafish and new mouse models, along with the associated kidney phenotypes.

## Results

### *Identification of compounds able to rescue* NPHP1-*associated cellular phenotypes*

We designed an *in vitro* high-throughput drug-screen strategy based on the Prestwick Chemical Library™, a collection of 1120 off-patent small molecules of which 95% are approved by FDA, EMA or other agencies. The screen was performed on the two most widely used renal cell models to study ciliogenesis and ciliopathies, i.e. the mouse inner medullary collecting duct (mIMCD3) and canine Madin-Darby Canine Kidney (MDCK) cells, in which the expression of *Nphp1* was knockdown through stable expression of shRNA (NPHP1_KD).

NPHP1_KD mIMCD3 cells display impaired ciliogenesis with a decreased percentage of ciliated cells, as we previously reported for NPHP1_KD MDCK cells (8), as well as increased cell migration during wound-healing assays (Sup Fig.1A-F). Using fluorescence microscopy-based automated strategies (Sup Fig.1G), 51 hits were identified based on their ability to positively modulate migration and/or cilium count in either mIMCD3 or MDCK NPHP1_KD cells. Hence, 11 compounds belonging to the different pharmacological classes were selected based on their effects on the analyzed phenotypes, on the absence of obvious toxicity (decrease cell count and/or inhibition of cell migration) and on their chemical action or physicochemical properties that may be relevant in NPH: known or suspected effects on cilium-associated pathways, actin cytoskeleton organization and/or microtubule polymerization, RhoGTPase activity, cAMP regulation, signaling pathways (e.g JNK, ERK/MAPK or Wnt) or their involvement in process such as fibrosis-related pathways or nephroprotective properties (Sup Fig.1G, Sup Table 1).

### Rescue of defective ciliogenesis in NPHP1 patients’ renal cells by Alprostadil, a PGE1 analogue

The efficiency of the 11 most interesting hits was further validated in urine-derived renal epithelial cells (URECs), a valuable source of cells from NPH individuals, previously described as a model to study ciliogenesis and ciliary functions in the context of ciliopathies (20). Primary URECs were collected either from individuals carrying bi-allelic mutations in *NPHP1* (NPHP1, hereafter Pt1-11), from age-matched control individuals (CTL, hereafter CTL1-5) or from individuals presenting with a non-ciliopathy-related chronic kidney disease (CKD, hereafter CKD1-4; Sup Table 2) and they were immortalized through the expression of a thermosensitive SV40 T-antigen. Transcriptomic analyses (Sup Fig.2A) showed that both primary and immortalized URECs exhibit similar expression levels of renal tubular epithelial cell markers, confirming their tubular origin that remained unaffected by the immortalization process. As expected, the immortalized URECs showed persistent proliferation at permissive temperature (33°C) but not at non-permissive temperature (39°C), as monitored by the loss of Ki67 and SV40 T-antigen nuclear staining in the latter condition (Sup Fig.2B-D).

In order to evaluate ciliogenesis in immortalized URECs, we set up a semi-automated pipeline based on basal-body (γ-Tubulin, red) and ciliary membrane (ARL13B, green) markers to quantify cilium count (Fig.1A). At non-permissive temperature, URECs from *NPHP1* individuals presented a significant decrease of cilium count compared to controls (Fig.1B, Sup Table 2). This ciliogenesis defect was associated with an increased proportion of cells presenting with two spots of CP110 per centrosome, indicative of an early blockage of ciliogenesis in NPHP1 URECs (Sup Fig.3).

**Figure 1:**
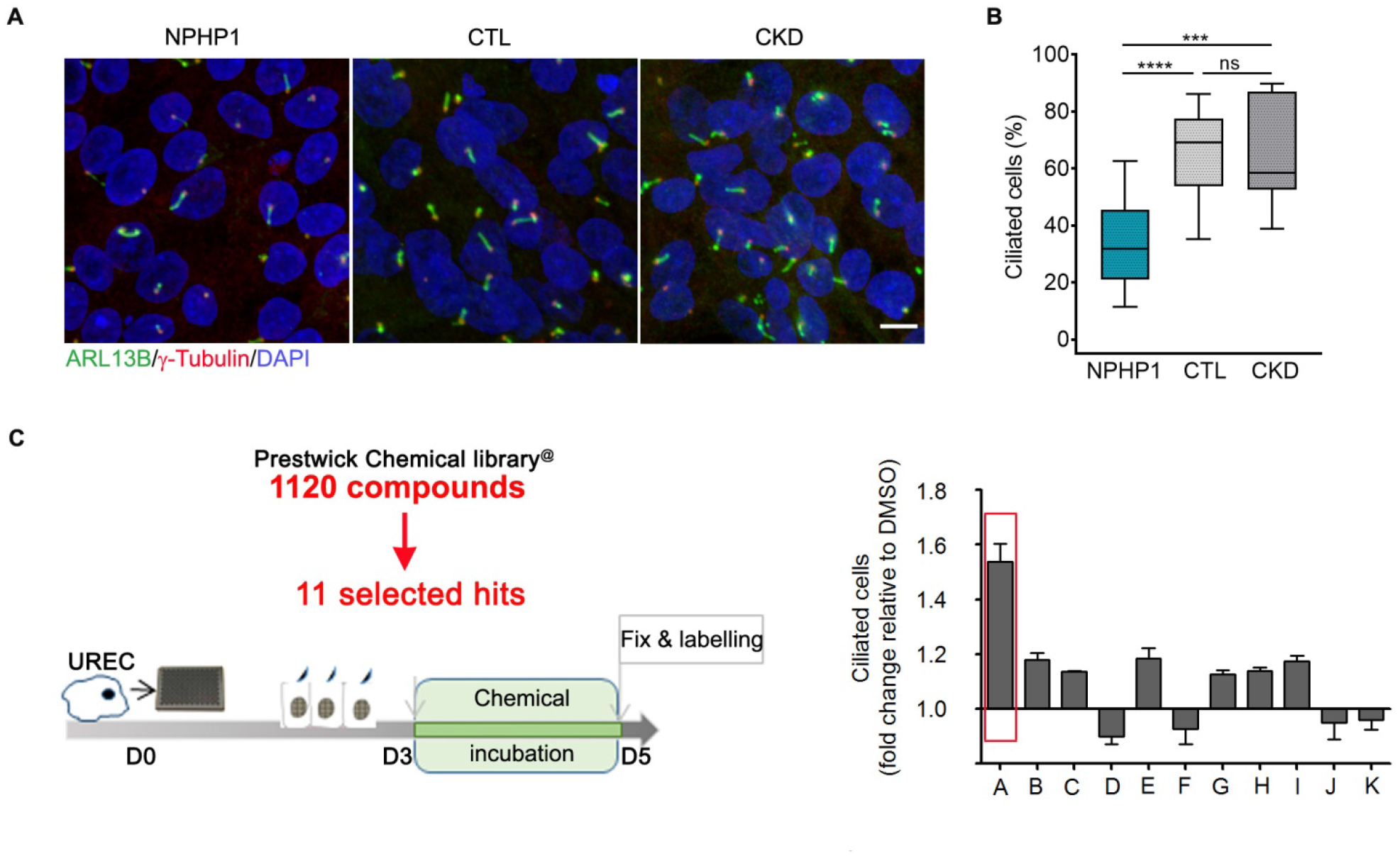
Drug Screens in NPHP1 defective renal cells. **(A)** Representative images of immunofluorescence of primary cilia (ARL13B, green), basal bodies (γ-Tubulin, red) and nuclei (DAPI, blue) in immortalized URECs derived from NPHP1 patients (NPHP1), control individuals (CTL) and CKD patients (CKD). Scale bar: 10μM. **(B)** Quantification of ciliogenesis in immortalized NPHP1, (*n* = 11), CTL (*n* = 6) and CKD (*n* = 5) URECs. Results are expressed as the mean for all individuals category and are shown using a box-and-whisker plot. *n* = 1–15 experiments, with a mean of 3.5 experiments per cell line Mixed linear regression model with quasibinomial penalization taking into account the correlation of observations coming from the same individuals using a random effect on the cell line: \*\*\**P* < 0.001, \*\*\*\**P* < 0.0001, ns: not significant. **(C)** Left panel: Immortalized NPHP1, CTL and CKD URECs were treated for 2 days with the 11 selected from the primary screen compounds and ciliogenesis was evaluated 5 days after seeding. Right panel: Quantification of ciliogenesis in one immortalized NPHP1 UREC line (Pt1) exposed to the 11 selected compounds. A: Alprostadil, 200nM – B: Cyproheptadine, 1μM – C: Ethopropazine, 50nM – D: Fluticasone, 10nM – E: Methotrexate, 400nM – F: Mycophenolic acid, 200nM – G: Paclitaxel, 100pM – H: Pyrimethamine, 500nM – I: Simvastatin, 10nM – J: Tropisetron, 2μM – K: Verapamil, 80nM. Mean ± SEM. *n* = 1–2 experiments.

Before assessing the effect of the 11 molecules on URECs, we evaluated the non-toxic range of doses at which they could be used (Sup Table 1). The highest non-toxic concentration as well as 1-log and 2-log concentrations were chosen to carry out the ciliogenesis assay on one of the *NPHP1* UREC line (Pt1).

Among the 11 molecules, a robust increase of cilium count was observed for Alprostadil (+1.53), a synthetic analog of prostaglandin E_1_ (PGE1), whereas the other compounds showed modest positive or negative effects on ciliogenesis (Fig. 1C). PGE1 (Alprostadil) is an analogue of the eicosanoid PGE2 which exerts various effects in inflammation, pain, vascular tone or renal function which are mediated by its receptors EP1-4 (encoded by *PTGER1-4*) (21). Notably, EP1-4 receptors are expressed in the URECs as revealed by RT-qPCR analyses (Sup Fig. 4A).

We first performed a dose-response analysis for Alprostadil (ALP) and determined the half maximal effective concentration (EC_50_) at 13.75nM while the maximal effect on cilium count was observed for concentrations above 1μM (Fig.2A, B). Based on these results, following experiments were performed using 2μM Alprostadil. In addition, Dinoprostone (DINO), a synthetic version of PGE2, showed a similar effect to Alprostadil (Sup Fig.4B), supporting the positive impact of prostanoids on cilia count in NPHP1 URECs. The positive effect of Alprostadil on ciliogenesis was confirmed in other NPHP1 URECs (Pt3-4,6-11; Fig.2C, Sup Fig.4C, D) but not in CTL or CKD cells, even in the one with a low proportion of ciliated cells (Sup Fig.4C, D), indicating the specific effect of Alprostadil on NPHP1 URECs. Furthermore, beyond the restoration of ciliogenesis, Alprostadil also restored ciliary composition with an increased percentage of Adenylate Cyclase 3 (ADCY3)-positive cilia, associated with an increased intensity of ciliary ADCY3 (Fig.2D-F).

**Figure 2:**
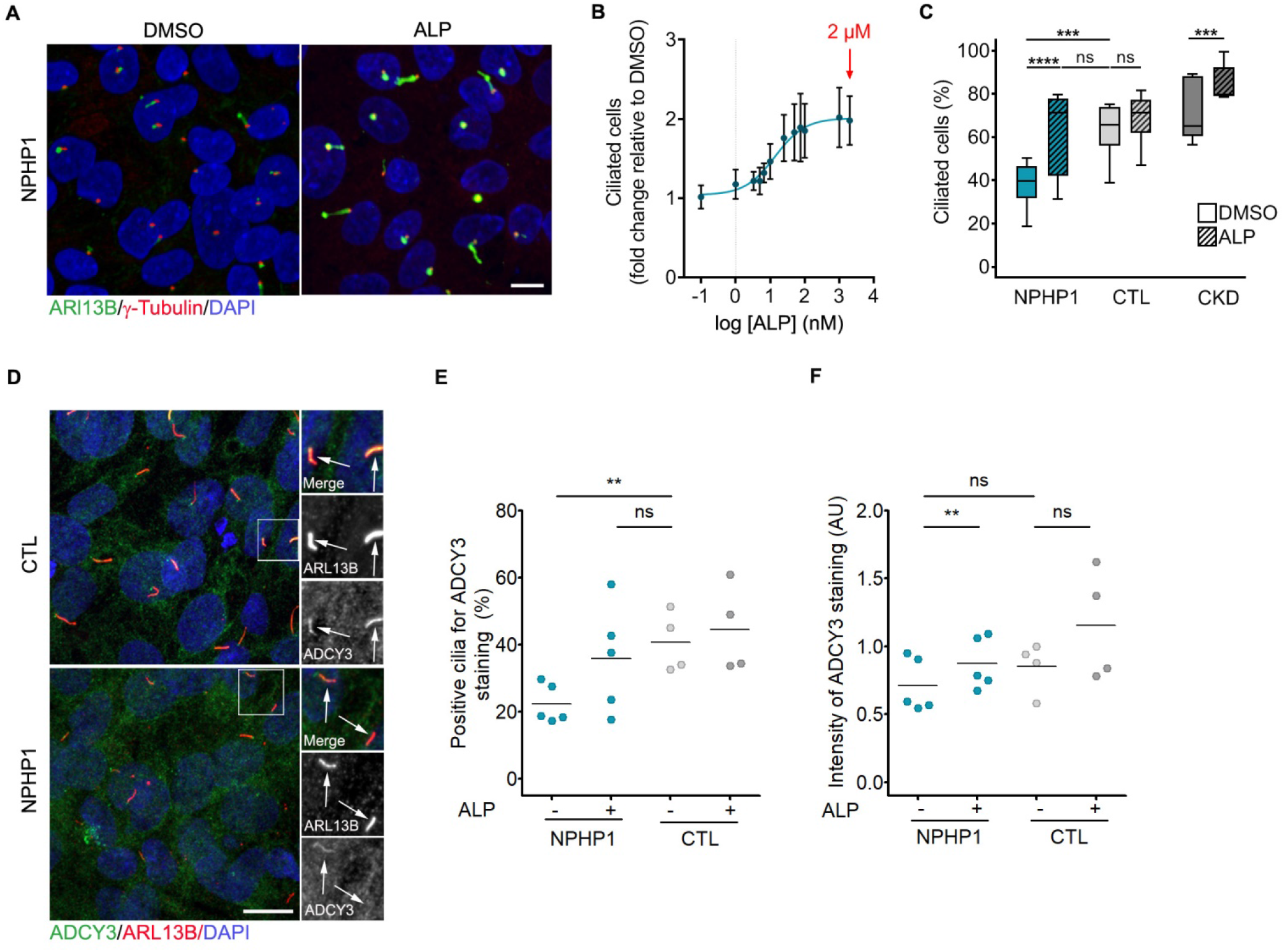
Validation of prostaglandin signaling as a target in *NPHP1* defective renal cells. **(A)** Representative images of immunofluorescence of primary cilia (ARL13B, green), basal bodies (γ-Tubulin, red) and nuclei (DAPI, blue) in immortalized NPHP1 URECs 5 days after seeding treated for 2 days with DMSO (0.04%) or Alprostadil (2μM). **(B)** Quantification of ciliogenesis in one immortalized NPHP1 UREC line (Pt1) exposed for 2 days to increasing concentrations of Alprostadil. Bars indicate mean ± SD. *n* = 1–4 experiments. Nonlinear regression. **(C)** Quantification of ciliogenesis in immortalized URECs derived from NPHP1 patients (NPHP1, *n* = 11), control individuals (CTL, *n* = 6) and CKD patients (CKD, *n* = 5) exposed for 2 days to DMSO (0.04%) and Alprostadil (2μM). Results are expressed as the mean for all individuals category and are shown using a box-and-whisker plot. *n* = 1–15 experiments, with a mean of 3.5 experiments per cell line. Mixed linear regression model with quasibinomial penalization taking into account the correlation of observations coming from the same individuals using a random effect on the cell line: \*\*\**P* < 0.001, \*\*\*\**P* < 0.0001, ns: not significant. **(D)** Representative images of immunofluorescence of ADCY3 (green), primary cilia (ARL13B, red) and nuclei (DAPI, blue) in immortalized NPHP1 and CTL URECs. **(E, F)** Quantification of the percentage of ADCY3 positive cilia (E) and the intensity of ADCY3 in those cilia (F) in immortalized NPHP1 (*n* = 5) and CTL (*n* = 4) URECs. Each dot represents one individual cell line. Bars indicate mean. *n* = 3 experiments. Mixed linear regression model with quasibinomial penalization (E), Mann-Whitney test (NPHP1 vs CTL) and paired Student t-test (DMSO vs ALP) (F): \*\**P* < 0.01, ns: not significant. AU: arbitrary unit. **(A, D)** Scale bar: 10μM. ALP: Alprostadil.

### In vivo validation of prostaglandin-related molecule in murine Nphp1 knockout model and NPH zebrafish model

We next aimed to test the therapeutic potential of PGE1 in a *Nphp1* mouse model. Using the CRISPR-Cas9 technology, we generated a *Nphp1* null allele which harbors a deletion of the start codon within the first exon of *Nphp1* (Sup Fig.5), resulting in a complete loss of *Nphp1* mRNA and protein expression, similarly as observed in most *NPHP1* patients (22). The *Nphp1^-/-^* mice displayed a severe retinopathy (Sup Fig.6) as described previously (23, 24). Histological examination of the kidneys revealed the presence of tubular dilatations in the cortex of *Nphp1^-/-^* mice as early as 2 months of age and more evidently at 5 months of age (Sup Fig.7A-E). A strong association was observed between the number and size of dilatations per kidney, and the “dilatation index” values for these two parameters showed a progression of kidney damages between 2 and 5 months in KOs compared with control animals (Sup Fig.7F-I).

Further characterization of these dilatations revealed that they mainly affect the connecting tubules (CNT) as well as the junction between the CNT and the distal convoluted tubules (DCT2) (Sup Fig. 8A, B). Notably, in kidney biopsies from patients carrying homozygous deletion of *NPHP1*, tubular dilatations were observed in similar tubular segments as well as in the loop of Henlé (Sup. Fig.8C). These data pointed to a specific role for *NPHP1* in the homeostasis of the DCT/CNT region both in mice and humans. Despite the presence of tubular dilatations, renal function was not impaired in 5-month-old *Nphp1^-/-^* mice (Sup. Fig.7J-M), indicating an early stage of the disease, consistent with a juvenile form of the disease in humans.

We next investigated whether treatment with the PGE1 analog Alprostadil could have a positive effect on the number and size of tubular dilatations in 5 months-old *Nphp1^-/-^* mice. Alprostadil was administrated intraperitoneally (*i.p.* 80μg/kg) daily to 1-month-old male *Nphp1^+/+^* and *Nphp1^-/-^* mice for 4 months without any impact on growth or survival (Sup Fig.9A). Histological analyses revealed that the number and total area of tubular dilatations were reduced in *Nphp1^-/-^* mice by the Alprostadil treatment with a linear correlation (Fig.3A, B) and a significant decrease in the dilatation index (Fig.3C, D). In addition, measurements of cilium length showed that cilia were longer in the dilated regions of *Nphp1^-/-^* mouse (Fig.3E, F, Sup Fig.9B), as observed in other NPHP models (25). A similar observation was obtained in kidneys of patient with *NPHP1* mutations compared to controls (Sup Fig.8D, E). In contrast, cilia length was reduced in Alprostadil-treated *Nphp1^-/-^* mice with values similar to those in control mice (Fig.3E, F, Sup Fig.9B). Despite the absence of detectable fibrosis by histology, we performed qPCR analysis of several profibrogenic markers in the kidneys of 5 months old *Nphp1^-/-^* mice (Fig.3G, Sup Fig.9C-G). Expression of the pro-fibrotic marker *Col1a1* was slightly increased in the kidneys of *Nphp1* mice, which was attenuated by Alprostadil treatment (Fig.3G), suggesting its potential beneficial effect on the development of renal fibrosis.

**Figure 3:**
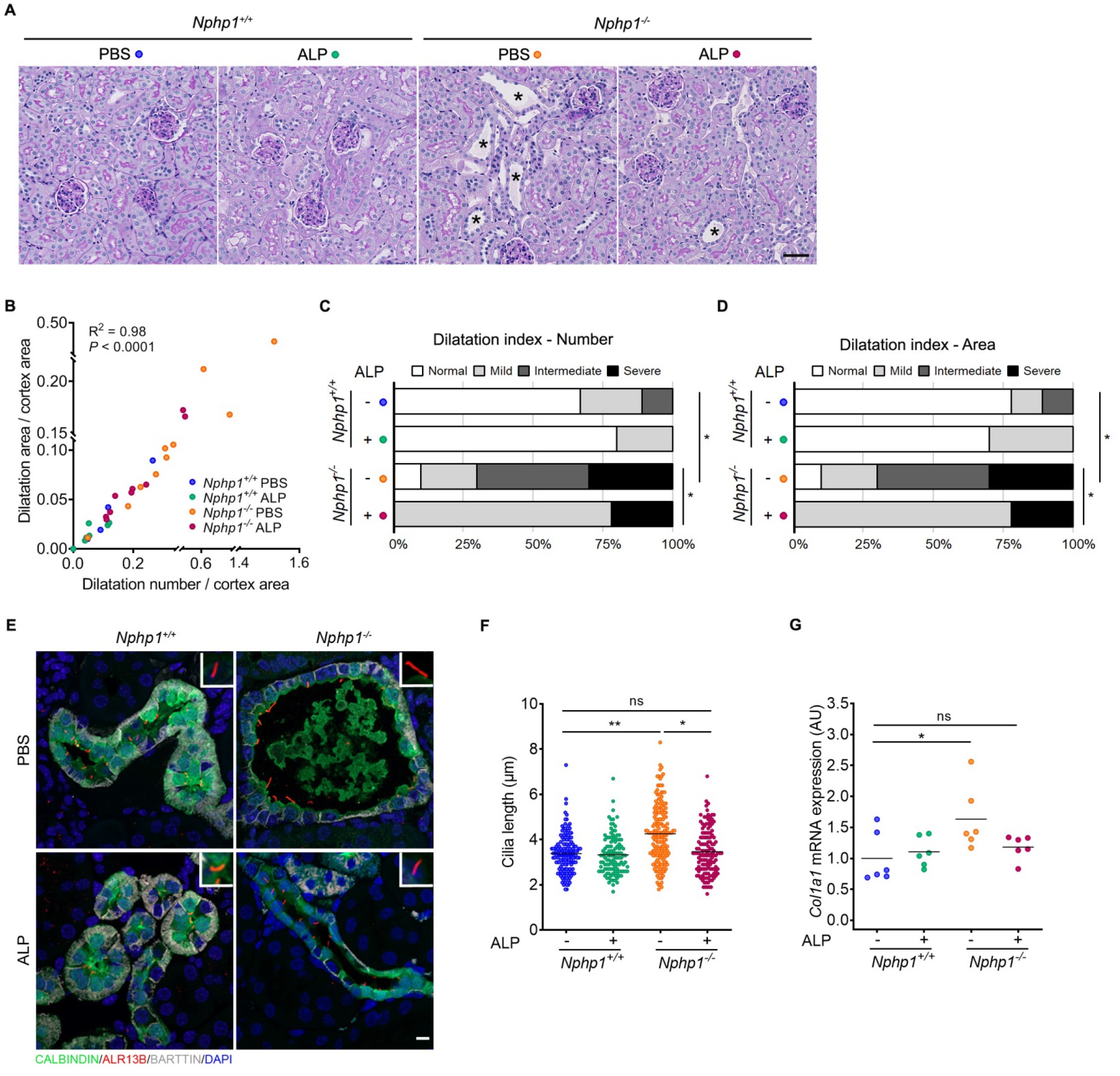
*In vivo* validation of the effect of Alprostadil treatment on *Nphp1^-/-^* renal tubule dilations. **(A)** Representative images of PAS staining of kidney sections from 5 months-old *Nphp1^+/+^* and *Nphp1^-/-^* mice treated daily with vehicle (PBS) or Alprostadil (80μg/kg) from 1 month to 5 months. Scale bar: 50μm. * = tubular dilatations. **(B)** Scatter dot plot correlating the number of dilatation/10μm^2^ in cortex to the total area covered by dilatations/cortex area in *Nphp1^+/+^* and *Nphp1*^-/-^ mice treated with PBS or Alprostadil (*n* = 9–10 male mice for each genotype/treatment). Pearson’s test (*r* = 0.9932). **(C, D)** Dilatation index for the number of dilatation/10μm^2^ in cortex (C) and the total area covered by dilatations/cortex area (D). Chi-squared test: \**P* < 0.05. **(E)** Representative images of immunostaining of 5 months-old *Nphp1^+/+^* and *Nphp1^-/-^* kidney sections upon Alprostadil treatment with anti-Calbindin (CNT, green), anti-ARL13B (cilia, red), anti-BARTTIN (TAL, DCT and CNT, gray) antibodies and DAPI (nuclei, blue). Scale bar: 100μm. **(F)** Quantification of cilium length (*n* = 3 male mice for each genotype/treatment). One-way ANOVA followed by Holm-Sidak’s post-test: \**P* < 0.05, \*\**P* < 0.01, ns: not significant. **(G)** RT-qPCR analysis of *Col1a1* expression from 5 months-old *Nphp1^+/+^* and *Nphp1^-/-^* kidneys upon Alprostadil treatment. One-way ANOVA followed by Holm-Sidak’s post-test: \**P* < 0.05, ns: not significant. AU: arbitrary unit. **(B, G)** Each dot represents one individual mouse. **(F, G)** Bars indicate mean. ALP: Alprostadil, TAL: Thick Ascending Limb of the Loop of Henle, DCT: Distal Convoluted Tubule, CNT: Connecting Tubule.

Overall, these data showed that Alprostadil treatment is able to attenuate the manifestations associated with renal ciliopathy, including the formation/extension of tubular dilatations, early upregulation of fibrosis markers and abnormal cilia.

Along with the mouse model, the effect of prostaglandin-related analogs was further examined in zebrafish, a widely used model to study cilia function in context of ciliopathies (26). We used the *nphp4* morpholino (MO) model, for which we and other have reported cysts in the proximal part of the pronephros as well as defects in cloaca formation and ciliogenesis in its distal end (Sup Fig. 10) (27, 28). This model is relevant as NPHP1 and NPHP4 play similar roles at the transition zone and cellular junctions; and their loss of function in patients as well as in other animal models results in identical phenotypes (1, 2). In addition, EP receptors in particular EP2 and EP4 are expressed in zebrafish pronephros (29). Treatments with both PGE2-derived molecules resulted in a significant decrease in the proportion of *nphp4*-morphants with severe pronephric cysts (Sup Fig.10A-C). Alprostadil treatment also reduced the proportion of *nphp4*-morphants with cloacal dilatations (Sup Fig.10D) and partially restored cilium length (Sup Fig.10E). Overall, these results obtained in zebrafish provide evidence that PGE2 analogs can prevent tubular dilatations and cilium defects in other *in vivo* models of NPH.

### Molecular analysis of Alprostadil treatment in URECs

To better understand the mechanism underlying the beneficial effect of PGE1 on ciliogenesis, we sought to determine which signaling pathways were involved downstream of EPs receptors in URECs. RT-qPCR analyses showed a predominant expression of *PTGER2* (EP2) and to a lesser extend *PTGER4* (EP4) compared with other EPs (Sup Fig. 4A), suggesting that the positive effect of Alprostadil was mostly mediated by EP2 and EP4. Notably, both of these receptors are coupled to GαS and have been shown to stimulate cAMP production through activation of adenylate cyclases (AC) in response to agonist stimulation (21). In addition, stimulation of cAMP production with the AC activator forskolin or EP4 agonists has been shown to exert positive effect on ciliogenesis (30). Accordingly, treatment with Alprostadil increased cAMP production in URECs to a level similar to that of Forskolin (Fig.4A) and increased the phosphorylation of protein kinase A (PKA) substrates (Fig.4B, C). Taken together, these results show that Alprostadil treatment activates the cAMP/PKA pathway, which may explain the improved ciliogenesis in *NPHP1* URECs. In agreement with this statement, forskolin treatment showed a similar positive effect on ciliogenesis as that observed with Alprostadil (Fig.4D, E).

**Figure 4:**
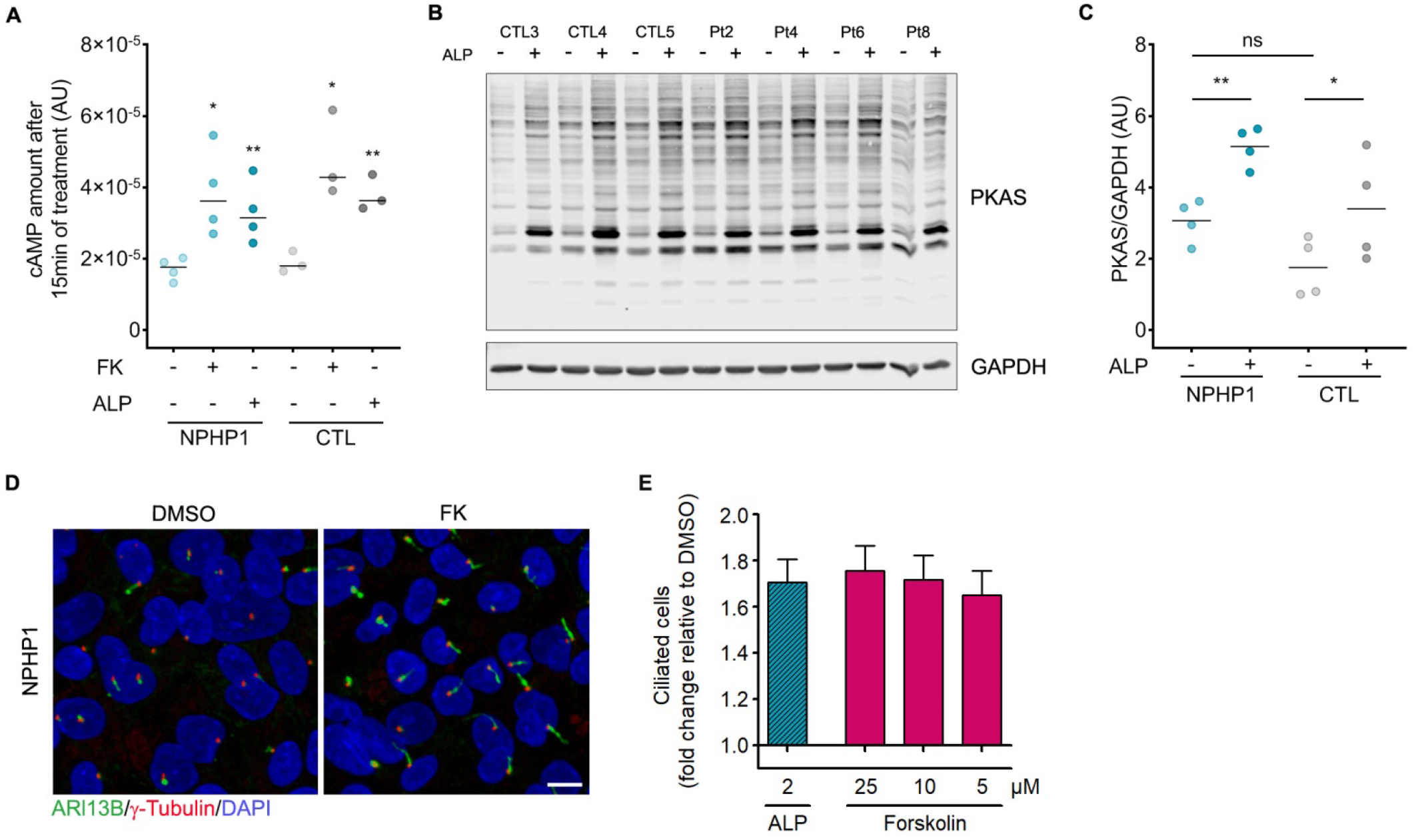
cAMP pathway upon Alprostadil treatment in NPHP1 URECs. **(A)** cAMP amount after 15 minutes of treatment with DMSO (0.04%), Forskolin (FK, 25μM) or Alprostadil (2μM) in immortalized URECs derived from NPHP1 patients (NPHP1, *n* = 4) and control individuals (CTL, *n* = 3). *n* = 3 experiments. Paired Student t-test (NPHP1/CTL DMSO vs NPHP1/CTL FK and ALP): \**P* < 0.05, \*\**P* < 0.01. **(B)** Representative image of Western blot analysis showing all phospho-PKA substrates using PKAS antibody in immortalized NPHP1 (Pt) and CTL URECs after 15 minutes of treatment with DMSO (0.04%) or Alprostadil (2μM). **(C)** Quantification of the relative abundance of PKAS in immortalized NPHP1 (*n* = 4) and CTL (*n* = 3) URECs. *n* = 3 experiments. Mann-Whitney test (NPHP1 vs CTL) and paired Student t-test (DMSO vs ALP): \**P* < 0.05, \*\**P* < 0.01, ns: not significant. **(D)** Representative images of immunofluorescence of primary cilia (ARL13B, green), basal bodies (γ-Tubulin, red) and nuclei (DAPI, blue) in immortalized NPHP1 URECs 5 days after seeding treated for 2 days with DMSO (0.04%) or Forskolin (25μM). Scale bar: 10μM. **(E)** Quantification of ciliogenesis in one immortalized NPHP1 UREC line (Pt6) exposed for 2 days to 2μM Alprostadil or increasing concentrations of Forskolin. Bars indicate mean ± SEM. *n* = 2 experiments. **(A, C)** Each dot represents one individual cell line. Bars indicate mean. AU: arbitrary unit. ALP: Alprostadil.

To identify functional downstream signaling pathways associated with the positive effects of PGE1 in NPHP1 models, we performed comparative transcriptomic profiling analysis in *NPHP1* URECs. First, we performed transcriptomics analysis in *NPHP1* (Pt1) URECs (Pt1) exposed to either DMSO or 2μM Alprostadil under the same conditions as for ciliogenesis assays and identified 457 differentially expressed protein-coding genes (150 upregulated, 307 downregulated). In parallel, we employed RNA-sequencing analysis on *NPHP1* URECs (including Pt2 and Pt8) and control lines (including CTL5 and CTL8), and revealed 2259 dysregulated genes in absence of *NPHP1.* We then compared the two gene lists and identified 128 dysregulated genes in *NPHP1* URECs (31 genes downregulated and 97 upregulated) that were inversely modulated upon Aprostadil treatment (Fig.5A, Sup Table 3). Pathway enrichment analysis using the GO, KEGG and Reactome datasets revealed that the 128 dysregulated genes were predominantly involved in the regulation of actin cytoskeleton (*P* = 1.00E-06), focal adhesion (*P* = 7.9E-06), extracellular matrix-receptor interaction (*P*=2.5E-06) and integrin cell surface interactions (*P* = 4.7E-05). In addition, several pathways crucial for mitochondrial function commonly impaired in chronic kidney diseases were also modulated, including positive regulators of fatty acid oxidation (FAO; *P* = 4.4E-04) (31) and the response to hypoxia (*P* = 9.1E-07) (32).

**Figure 5:**
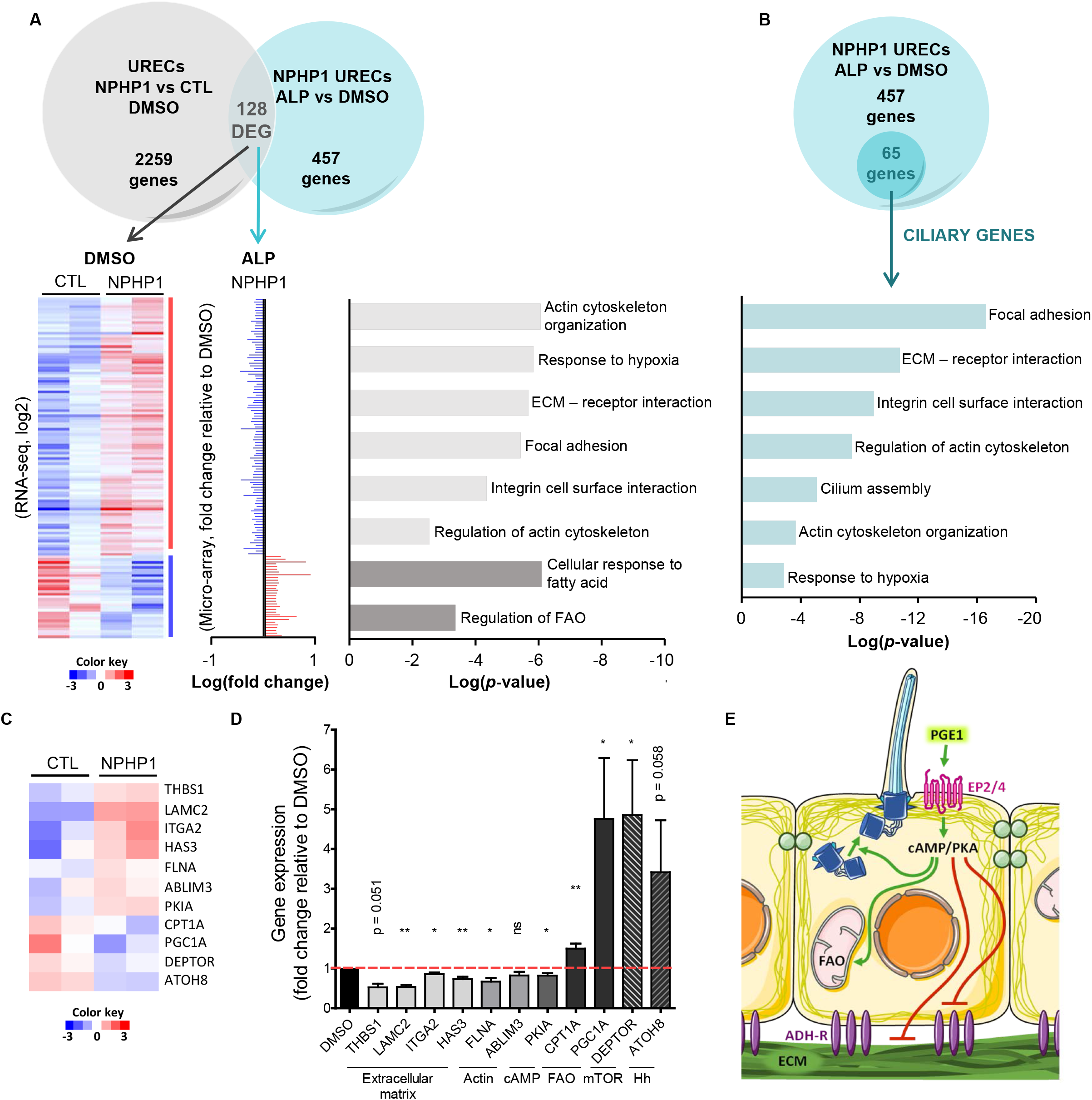
Transcriptomic analyses of molecular targets upon Alprostadil treatment in NPHP1 URECs. **(A)** Venn-diagram shows the number of significantly differentially expressed genes detected by microarray in immortalized URECs derived from NPHP1 patients (NPHP1) and control individuals (CTL) upon Alprostadil (ALP) treatment versus DMSO (457 regulated genes) in comparison to the number of deregulated genes detected by RNA-Seq in NPHP1 versus CTL URECs (2259 regulated transcripts). The 128 genes regulated by ALP inversely modulated in NPHP1 URECs (128 DEG) are underlined. Left panel: Heatmap of the 128 DEG genes from the RNA-Seq dataset of NPHP1 (*n* = 2) and CTL (*n* = 2) URECs using signal log ratio values. Each row represents a gene with up-regulated (red) expression and down-regulated (blue) expression. Middle panel: Fold change and normalized expression of the 128 DEG genes in one NPHP1 UREC (Pt1) selected from the microarray dataset. *n* = 3 experiments. Right panel: pertinent selected pathways or relevant processes involving these genes were highlighted using Metascape. **(B)** Diagram shows the number of ciliary modulators deregulated in NPHP1 URECs upon ALP treatment versus DMSO (65 genes). Pertinent selected pathways or relevant processes involving these genes were highlighted using Metascape. **(C)** Heatmap of 11 selected genes out of the 128 DEG genes from RNA-Seq dataset of NPHP1 versus CTL URECs. **(D)** RT-qPCR validating the positive effect of ALP treatment on the expression of the 11 selected deregulated genes in NPHP1 URECs (*n* = 5) (extracellular matrix-related genes; actin-related genes; cAMP signaling-related genes; fatty acid oxidation (FAO)-related genes; mTOR signaling-related gene; Hedgehog (Hh) signaling-related gene). Paired Student t-test: \*\**P* < 0.01, \**P* < 0.05, ns: not significant. Mean ± SEM. *n = 2* experiments. **(E)** Pathways regulated by Alprostadil and cAMP/PKA. PGE1: prostaglandin E1, EP2/4: prostaglandin E2 receptor 2/4, cAMP: cyclic adenosine monophosphate, PKA: protein kinase A, FAO: fatty acids oxidation, ADH-R: adhesion receptors, ECM: extracellular matrix.

In addition to the common-enriched pathways highlighted, several deregulated genes were also associated with renal fibrosis, including the renal profibrotic factors *GREM1* and *THBS1* (33, 34) as well as negative regulators of the mTOR pathway (*DEPTOR*) (35) or cyclic AMP signaling (*PDE7, PKIA, ANX3*), consistent with known signaling pathways downstream of EPs (36) and with our observations. Furthermore, when the 457 ALP-regulated genes were compared to a comprehensive list of genes encoding known ciliary proteins or ciliogenesis modulators, 65 were found to be cilium-related (Fig. 5B, Sup Table 4) – in agreement with the positive effect of Alprostadil on ciliogenesis.

To validate our transcriptomic analyses, we then selected 11 dysregulated genes belonging to the main pathways/cellular processes identified, including genes associated with cell adhesion (*LAMC2, ITGA2*), renal fibrosis (*THBS1, HAS3, LAMC2, ITGA2*), regulation of cAMP (*PKIA*), mTOR (*DEPTOR*) and FAO (*CPT1A, PGC1A*), as well as potential modulators of ciliogenesis (*ABLIM3, ATOH8*) several of which have already been shown to be protective against renal fibrosis and inflammatory processes (*CPT1A, PGC1A, DEPTOR*; Fig.5C). RT-qPCR analyses in several *NPHP1* URECs confirmed the effect of Alprostadil treatment on both the up- and down-regulation of most of the genes tested (Fig.5D). PGE1 treatment downregulated mRNA expression of all selected extracellular matrix components and integrin signaling regulators (*THBS1, HAS3, LAMC2, ITGA2).* In contrast, the expression of the two positive FAO regulators, *CPT1A* and *PGC1A*, and the inhibitor of mTOR signaling *DEPTOR*, was upregulated upon PGE1 treatment. Finally, according to their effect on ciliogenesis, *ABLIM3* was downregulated after PGE1 treatment, whereas the gene encoding the atonal bHLH transcription factor family member, *ATOH8*, was upregulated by Alprostadil treatment. This is coherent with *ATOH8* being involved in kidney development, Hh signaling, and may have a potential positive role on ciliogenesis as described for its homolog Atoh1 (37, 38).

Altogether, our results suggest that a prostaglandin compound would be able to restore ciliary and polarity defects in *NPHP1*-deficient cells through modulation of actin, adhesion and extra-cellular matrix cue (Fig.5E).

## Discussion

There are approximatively less than 5% of treatment available for the more than 7000 orphan diseases so far identified – most of which have genetic causes (39). The work presented here exemplifies the principle of identification of small pharmacological molecules to compensate genetic defects as source for potential therapeutic agents. A phenotypic screening strategy using a reference cell line derived from a *NPHP1* patient led to the rapid hit identification of a prostaglandin (PGE1, Alprostadil) that reverted whole-locus *NPHP1* deletion-associated ciliary phenotypes *in vitro* in a panel of *NPHP1* patients-derived cell lines. *In vivo*, PGE1 lessened the renal phenotypes observed in both a NPH zebrafish model and novel *Nphp1* knock-out murine model.

### Variability of the ciliary phenotype in NPHP1 models

Although ciliary defects observed in the different NPH *in vitro* and *in vivo* models are variable, they are all rescued by PGE1 treatment. Our present and past analyses consistently indicate that the loss of NPHP1 in kidney epithelial cells results in decreased ciliogenesis. This ciliogenesis defect corresponds to a decreased percentage of ciliated cells, which was linked to epithelialization delay in MDCK grown in 2D conditions (8). It is noteworthy that when NPHP1_KD MDCK cells are grown in 3D (spheroids), cilia get longer similarly as *in vivo*. Indeed, in *Nphp1* null mouse model and in patient kidney biopsies, cilia present at the apical surface of renal tubular cells are longer than in healthy individuals. As for the zebrafish NPH model, ciliogenesis is decreased and cilia are shorter in the dilated distal pronephros as observed in other ciliopathy fish models (26). Importantly, the length of the primary cilia depends on various factors, including, microtubule and actin cytoskeleton dynamics, signaling pathways such as ROS, mTOR, cAMP, cell cycle and cell epithelialization, and may be differentially altered in NPH genes/models with as a consequence either shortening or elongating of primary cilia (1, 39). This diversity of processes that governs ciliogenesis could explain why, depending of the cellular or animal models, we observed different ciliary phenotypes.

Thus, decreased ciliogenesis in *in vitro* NPHP1-associated phenotype in kidney cells grown in standard culture conditions is a robust proxy, which led to the identification of a molecule that restores ciliogenesis defects in all kinds of NPH models.

### Ciliogenesis and cAMP

Several studies support the principle of our findings linking EP receptors and ciliogenesis. It has been shown that defective levels of prostaglandin synthases, PTGES1 and PTGES2, result in the loss of connecting cilia and photoreceptor cell differentiation in zebrafish models (40). Additionally, stimulation of EP4 positively modulates ciliogenesis *in vitro*, as well as *in vivo* in zebrafish, through the production of cAMP and the activation of anterograde IFT (30). It is worthwhile noticing that ciliopathies associated with mutations in *NPHP* or *IFT* genes have been independently linked to a downregulation of cAMP/PKA signaling associated with a decrease of ciliary localization of ADCY3 (9, 10, 41). Finally, treatment by forskolin, an agonist of ACs, was shown to ameliorate cilium count and length in *CEP290* defective fibroblasts (10). In this study, we confirm that PGE1 treatment increase the percentage of ADCY3-positive cilia in *NPHP1* URECs, as well as the cAMP production and PKA activation. Moreover, treatment by forskolin rescue ciliogenesis in *NPHP1* URECs, confirming that the positive effect of PGE1 in those cells is likely mediated by cAMP production *via* the activation of EP2 and EP4 receptors.

In line with these reports, our transcriptomic analyses of PGE1-treated URECs identified several cAMP regulators, in particular the cAMP-Dependent Protein Kinase Inhibitor Alpha, PKIα (42). The upregulated expression of *PKIA* in the absence of *NPHP1* is expected to lead to decreased activity of PKA, which would be counterbalanced by activation of EP2 and EP4 receptors upon PGE1 treatment through the inhibition of PKIα and increased cAMP production. Dysregulation of cAMP signaling is a common feature of CKD. In ADPKD and infantile NPH mouse models, renal cAMP levels are increased and associated with increased proliferation and cyst formation (43, 44). In contrast, cAMP signaling cascade is decreased in several models of renal fibrosis (45). Whether cAMP signaling pathway is invovled in juvenile NPH is still unclear; however, we did not observed any increase of cAMP production in *NPHP1* URECs at a basal level, and juvenile NPH is mainly characterized by small kidneys with massive renal fibrosis and tubular dilatations, associated with apoptosis and absence of cell proliferation (45, 46). Thus, even if alterations of cilium-mediated mechanisms are involved in the pathogenesis of both juvenile NPH and ADPKD, it is likely that distinct signaling pathways and/or regulation levels are impacted in these two diseases.

### Ciliogenesis and cell architecture

Besides the putative direct positive impact of EP receptors-mediated cAMP signaling on ciliogenesis, other prostaglandin-dependent pathways or cellular processes are susceptible to also modulate ciliation. It is well documented that cell shape, contractility and extracellular matrix rigidity are major determinants of ciliogenesis, with increasing evidence that proteins with actin binding or cell adhesion functions can control ciliary dynamics (46, 47). Moreover, PGE2 is known to favor epithelial maturation in retinal pigment epithelial cells derived from iPS cells mutated in ciliopathy genes (48).

Notably, prostaglandins are known to regulate actin cytoskeleton remodeling essential for cell motility, and cell-cell and cell-matrix interactions *via* activation of cAMP-dependent PKA or small GTPases Rac1 and Rho (49–51). In agreement with this, our transcriptomic studies of PGE1 treated *NPHP1* URECs showed that the actin-cytoskeleton regulation and extracellular matrix organization were the most enriched GO terms – with a large portion of deregulated genes encoding regulators of actin cytoskeleton organization, cell-substrate adhesion, cell migration and components of the integrin pathway. In particular, silencing of several integrin and ECM structural components altered cytoskeletal organization, enhanced ciliation and restrict cell migration similarly to PGE1 treatment in *NPHP1* invalidated cells. For that matter, cell migration was the initial phenotype restored in *Nphp1*_KD mIMCD3 by PGE1 treatment. Globally, the actin regulators, ECM components and integrins, overexpressed in *NPHP1* URECs, were lessened upon PGE1 treatment.

### Fibrosis

Renal fibrosis as seen in patients is rarely observed in murine NPHP models - except in the case of *Nphp7/Glis2* (52) or in a recent model mimicking NPH by invalidation of *Lkb1* in distal tubules (53). Although no interstitial fibrosis was visible in 5 months-old *Nphp1^-/-^* kidneys, we were able to detect an increased expression of profibrotic markers in *NPHP1* URECs or in *Nphp1^-/-^* kidneys, in particular the collagens *Col1a1* and *Col3a1* as well as their receptor, the integrin *ITGA2.* Notably, *Itga2* is particularly enriched in the thick ascending loop of Henle, the distal and the connecting tubules (54) which are the nephron segments altered in *Nphp1* KO mice and patients. This is coherent with an early description of a specific increased expression of ITGA2 in NPH patient biopsies (55) and with the remodeling and thickening of the basement membrane observed in NPH.

In addition, transcriptomic analysis in URECs pointed to several other profibrotic genes including *THSB1*, encoding Thrombospondin 1, a extracellular matrix protein that has diverse roles in regulating cellular processes important for the pathogenesis of fibrotic diseases ^33^ and *GREM1*, a negative regulator of BMP signaling which was shown to directly participate in renal fibrosis in CKD models (33). Treatment with PGE1 tends to reduce *THSB1, LAMC2, ITGA2* and *GREM1* expression in *NPHP1* URECs, suggesting a potential positive effect of these compounds on renal fibrosis. In accordance, EP2 and/or EP4 activation exert anti-fibrotic properties, as observed in several *in vitro* and *in vivo* models of kidney fibrosis (56, 57). These results suggest that the *NPHP1* deficient background (KO mice and URECs), in contrast to the control background, is sensitive to PGE1 treatment, which has a protective effect on profibrotic gene expression and renal lesions.

### Fatty-acid oxidation (FAO) pathway

Beside the cell-adhesion and ECM enriched pathways, we highlighted the regulation of fatty-acid oxidation (FAO) as a novel pathway affected in the absence of *NPHP1*, and its relationship with PGE2/EP receptor signaling. FAO pathway is indispensable for renal tubular cells homeostasis, and defective FAO in renal tubular epithelial cells has a key role in kidney fibrosis development (31, 58). The two FAO components affected in *NPHP1* URECs*, CPT1A* and *PGC1A*, are predominantly expressed in the distal tubules in mammalian kidneys where mitochondrial activity also plays an important physiological function in DCT and CNT cells (59). The fact that the distal tube is the seat of lesions observed in NPH could suggest a specific role of the FAO pathway and metabolic processes in the appearance of lesions linked to the absence of *NPHP1.* Importantly, FAO signaling can be activated by the cAMP/PKA/CREB pathway in several tissues (59), thus explaining the positive activation of this pathway by the prostaglandin-mediated EP2 or EP4 treatment in *NPHP1* deficient cells.

### Conclusion

The action of PGE1/EP receptors signaling acts both on ciliogenesis and epithelial cell architecture: it is likely that ciliogenesis is restored through multiple mechanisms, which includes cAMP – and that cell architecture is possibly restored through possibly direct functions of several NPHP components in the cell-cell/cell-matrix protein complexes and microtubule/actin networks (60–63).

This study provides the first evidence for a potential use of a small molecule-based therapy in the specific context of the *NPHP1* whole-locus loss-associated NPH, the most frequent form of NPH. PGE2 analogues showed restoration effects on ciliary and renal pre-clinical NPHP1-associated manifestations, therefore providing interesting new potential therapeutic targets. More importantly, our study demonstrated that phenotype changes induced by a complete locus deletion are amenable to be reversed, and they can be functionally complemented with an indirect therapeutic molecular point of intervention using small molecules.

## Materials and Methods

### Human urine-derived renal epithelial cells (URECs)

Patients and control individuals were recruited at Necker Hospital in the frame of the approved NPH_1 protocol and urine samples were collected upon written informed consent and anonymized together with personalized data. URECs were isolated from urine collected from patients suffering from nephronophthisis carrying bi-allelic *NPHP1* mutations, healthy age-matched donors and patients suffered from ciliopathy-unrelated chronic kidney disease (CKD stage 2 to 5; Sup Table 2). URECs were cultured as previously described (63) with the following modifications. Briefly, peeled renal tubular epithelial cells were cultured in Primary medium (DMEM/F12, 10% FBS, 100U/mL penicillin, 100mg/mL streptomycin, amphotericinB, 1×REGM SingleQuots (CC-4127, Lonza)) in 6-well plates. Cells were incubated at 37°C in 5% CO2, and Primary UREC culture medium (0.5mL) was renewed every 24h. At 96h, the culture medium was replaced by Proliferation UREC medium REBM Basal Medium (CC-3191, Lonza) containing 1×REGM SingleQuots, 10ng/mL rhEGF (R&D system), 2% FBS certified (Invitrogen) and changed every 2 days. At 80% confluence, cells were trypsinized (TrypsinE select, Invitrogen) and plated for immortalization (2×10^4^ cells in 12-well plates). After 24h, cells were transduced by thermosensitive retroviral gene transfer of SV40 T antigen (LOX-CW-CRE, Addgene) at a MOI of 5 in Proliferation UREC medium with 8μg/mL polybrene (64). After 24h, the infection medium was replaced by fresh complete REGM, and cell culture was carried out at the permissive temperature of 33°C. After 3 passages, the SV40 T antigen expression was assessed by immunofluorescence at both 33°C and 39°C.

### Antibodies

The following antibodies were used in the study with the following dilution: ADCY3 (1/100^e^; Invitrogen, PA5-35382), ARL13B (1/800^e^; ProteinTech, 17711-1-AP), Barttin (1/400^e^; Santa Cruz, sc365161), Calbindin D-28K (1/500^e^; Novus, NBP2-50028SS), GAPDH (1/1000^e^; Millipore, MAB374), γ-Tubulin (1/5000^e^; Sigma, T6557), PKAS (1/1000^e^; Cell Signaling, 9624), DAPI (1/2000^e^; Thermo Fisher Scientific, 62247).

### Secondary Screen of the 11 selected hits on URECs

#### Cell culture, compound treatment and immunofluorescence

Patient and control URECs (10^5^cells/well) were seeded in 96-well glass microplates (Sensoplates, Greiner) at 39°C. On the third day of culture, cells were incubated for 48h in the presence of the drug. For each drug, 3 doses were tested, depending on the live-cell cytotoxicity assay. All drugs were dissolved in DMSO (0.04% to 0.1%). Alprostadil was diluted in 0.04% DMSO. Cells were fixed after 5 days of culture in cold methanol for 5 minutes, then treated 40 minutes with PBS – 0.1% Tween 20 – 1% BSA, before incubating 1h with primary antibodies, followed by washes and incubation with appropriate fluorescent secondary antibody.

#### Data acquisition, quantification, and analysis

Images were acquired within 3 days using the Opera Phenix (40x, Perkin Elmer). Automated acquisition of 41 Z-stack per well was performed. The number of ciliated cells was measured using a semi-automated analysis using Harmony software (Perkin Elmer). Briefly, images were analyzed using the building blocks approach to detect in this order, nuclei (DAPI staining) and, with a 20px enlarged region, the basal body (γ-Tubulin signal) and the primary cilia (ARL13B staining). With conventional filters (intensity, size...), the software segmented candidate primary cilium. Hundreds of phenotypic parameters were calculated for every candidate cilium using SER texture (intensity patterns) and advanced STAR morphology parameters (distribution of either texture features or fluorescence intensities inside a region of interest). Using the PhenoLOGIC™ machine-learning option of Harmony, the parameters best suited to discriminate cilia were defined and used to obtain final detection and counting of primary cilia. Because of high cell confluence, Harmony software failed to properly segment nuclei, thus the number of cells was based on basal bodies.

### Immunofluorescence on URECs

Cells were fixed after 1 to 5 days of culture in cold methanol for 5 minutes (SV40 and CP110 staining) or with 4% PFA for 20 minutes (ADCY3 staining), quenched in 50mM NH_4_Cl and permeabilized 15 minutes with 0.1% Triton if fixed with 4% PFA, then treated 30 minutes with PBS – 0.1% Tween 20 – 3% BSA, before incubating 1h with primary antibodies, and then with appropriate fluorescent secondary antibody. Confocal images were acquired using a Spinning Disk microscope (40x or 63x, Zeiss). All quantifications were performed using Image J software.

### cAMP dosage in URECs

Cells were incubated during 15 minutes with 0.04% DMSO, 25μM Forskolin or 2μM Alprostadil. cAMP production was measured using cAMP-Glo™ Assay (V1501, Promega) following the recommendations of the manufacturer.

### RNA extraction for transcriptomic analyses

URECs were seeded in condition favoring ciliogenesis: primary cells were seeded (2×10^4^ cells/well) in 2 wells of 12-well plates and incubated one week at 37°C, immortalized patients and controls cells were seeded (10^5^ cells/well) in 8 wells of 96-well glass microplates at 39°C and incubated for 4 days. To evaluate the impact of Alprostadil treatment on transcriptome, immortalized Pt1 UREC line was incubated for 24h on the third day of culture with DMSO or 2μM of Alprostadil. Total mRNA was isolated using Extraction Mini Kit (Qiagen) following the recommendations of the manufacturer.

### Microarray data analysis

Affymetrix Human ClariomD data were normalized using quantile normalization with adjustment based on the median intensity of probes with similar GC content (using Affymetrix Power Tools). Background correction was made using the antigenomic probes. Only probes targeting exons annotated from FAST DB v2016_1 transcripts were selected. Probes were considered as expressed if the DABG *P* value ≤ 0.05 (Detection Above Background *P* values were calculated using Affymetrix Power Tools) in more or equal than 60% of samples in at least one of the two compared experimental condition (*e.g.* RLT 10μM). Genes were considered as expressed if more or equal than 50% of their probes are expressed. Minimum 4 selected probes were required to assess gene expression. When possible, only high-specific probes were selected (*i.e.* not overlapping with repeat regions; not cross-hybridizing; and 40 ≤ GC% ≤ 60). In addition, if possible, only probes targeting constitutive gene regions were selected (*i.e.* targeting at least 75% of transcripts of a given gene). We performed a paired Student’s t-test to compare gene intensities in the different biological replicates. Genes were considered significantly regulated when fold change was ≥ 1.5 and uncorrected *P* value ≤ 0.05.

#### Unsupervised analysis

The PCA has been performed using “prcomp” function in R and the 2 first dimensions were plotted. The clustering has been performed using “dist” and “hclust” functions in R, using Euclidean distance and Ward agglomeration method. Bootstraps have been realized using “pvclust” package in R, with the same distance and agglomeration method, using 1000 bootstraps.

#### Sample reproducibility study

Pearson correlation tests have been performed for each pair of samples using “cor.test” function in R. Heatmaps and clusterings were performed with “dist” and “hclust” functions in R using Euclidean distance and Ward agglomeration method. Bootstraps were realized as described in “Unsupervised analysis” method. Analysis for enriched GO terms, KEGG pathways and REACTOME pathways were performed using the automated meta-analysis tool Metascape (http://metascape.org/).

### RNAseq analysis

For each sample, 50 million reads pair-end sequencing was performed on NovaSeq 6000 Illumina. FASTQ files were mapped to the ENSEMBL [Human(GRCh38/hg38) / Mouse GRCm38/mm10] reference using Hisat2 and counted by featureCounts from the Subread R package (http://www.r-project.org/). Read count normalizations and group comparisons were performed by the Deseq2 statistical method. Flags were computed from counts normalized to the mean coverage. All normalized counts < 20 were considered as background (flag 0) and ≥ 20 as signal (flag=1). P50 lists used for the statistical analysis regroup the genes showing flag=1 for at least half of the compared samples. The results were filtered at *P* value ≤ 0.05 and fold change 1.2. Cluster analysis was performed by hierarchical clustering using the Spearman correlation similarity measure and ward linkage algorithm. Heatmaps were made with the R package ctc: Cluster and Tree Conversion and imaged by Java Treeview software (Java Treeview – extensible visualization of microarray data) (65). Functional analyses were carried out using Metascape (http://metascape.org/).

### RT-qPCR analysis

Total RNA was reverse transcribed using SuperScript II Reverse Transcriptase (LifeTechnologies) according to the manufacturer’s protocol. Quantitative PCR was performed with iTaq Universal SYBR Green Supermix (Bio-Rad) on the ViiA 7 Real-Time PCR System (Thermo Fisher Scientific). Each biological replicate was measured in technical duplicates. The primers used for quantitative real-time PCR are listed in Sup Table 5.

### Mice experiments

#### Generation of Nphp1^-/-^ mice by CRISPR/Cas9

*Nphp1*^-/-^ mice were generated using a CRISPR/Cas9 system as previously described (66). Guide RNAs targeting exon1 of the gene were designed using CRISPOR (http://crispor.tefor.net/). sgRNA and Cas9 protein (TACGEN) were injected into the pronucleus of the C57Bl/6J zygotes. All exonic off-targets were tested to ascertain absence of undesired mutations. Two founders carrying indel mutations were backcrossed with C57BL/6j mice and the *Nphp1*^-/-^ offspring were further confirmed by PCR genotyping with appropriate primers. We focused on the mouse line carrying a 73bp deletion (c.-33_40del) in exon1, which deleted the ATG.

#### Histology and immunohistochemistry

Kidneys and eyes were fixed overnight at 4°C in 4% PFA and embedded in paraffin. The retinal and kidney structures were assessed by Hematoxylin/Eosin and Periodic Acid Schiff (PAS) staining of tissue sections (4μm), respectively. Count and area of kidney dilatations were measured using Image J software. For immunostaining, after paraffin removal and antigen retrieval treatment (10mM Tris pH9 – 1mM EDTA – 0.05% Tween 20, 87°C, 50 minutes), kidney sections were blocked with PBS – 0.1% Tween 20 – 1% BSA for 1h and incubated overnight at 4°C with appropriate primary antibodies. Cilia length was measured using Image J software. Confocal images were acquired using a Spinning Disk microscope (40x or 63x, Zeiss).

#### Dilatation index

In order to quantify more accurately the severity of the disease, a dilatation index, taking into account the dilatation number and dilatation size was established. For each calculated ratios (a) the dilatation surface area to cortex area and (b) the dilatation number to cortex area, a score was defined as following: score = 1 (no lesions) for all values between zero and the 3^rd^ quartile of all values for *Nphp1^+/+^*; score = 2 (mild) for all values between score 1 and the median value for *Nphp1^-/-^*; score = 3 (intermediate) for all values between score 2 and the 3^rd^ quartile of all values for *Nphp1^-/-^*; score = 4 (severe) for all values above score 3. Scores are protocol specific and calculated based on all the animals included in the protocol.

### Western blot analysis

Kidneys were lysed in 50mM Tris HCl pH8 – 200mM NaCl – 1mM EDTA pH8 – 1mM EGTA – 1mM DTT – 1% SDS. Cells were lysed in 50mM Tris HCl pH7.4 – 150mM NaCl – 0.1% SDS – 1% Triton – 0.5% sodium deoxycholate. Lysis buffer was supplemented with protease and protein phosphatase inhibitor tablets (04-693-159-001 and 04-906-845-001 respectively, Roche). Protein content was determined with the BCA Protein Assay kit (23225, Thermo Fisher Scientific). Equal amounts of protein were resolved on gels under reducing conditions, transferred and incubated with primary and secondary antibodies, and visualized using the Fusion Fx7 (Vilber) imaging system or the Odyssey ® CLx imaging system (LI-COR Biosciences).

### Statistics

Statistical calculations were performed with R or GraphPad Prism version 8. Figure representation was performed using GraphPad Prism version 8 and is presented as dot plots or box-and-whisker plots. The Shapiro-Wilk test was used to test the normality. Differences between 2 groups that do not meet the normal distribution were compared using two-tailed Mann-Whitney test. If not possible or if the 2 groups meet the normal distribution, differences were compared using unpaired two-tailed Student *t*-test. When comparing more than two groups a one-way ANOVA followed by Holm-Sidak’s multiple comparisons test was used. To analyze the effect of Alprostadil treatment on ciliogenesis or ADCY3 positive cilia staining in cells coming from the same individuals, we used a mixed linear regression model with quasibinomial penalization (R software: qlmmPQL function of MASS package) taking into account the correlation of observations coming from the same individuals using a random effect on the cell line. To analyze the effect of Alprostadil treatment in cells coming from the same individuals, we used a paired two-tailed Student *t*-test. To compare the correlation between two parameters, we used a Pearson’s test. To determine if there is a significant difference between two proportions, we used Fisher’s exact test. To determine if there is a significant difference between more than two proportions, we used a Chi-squared test. A *P* value of less than 0.05 was considered statistically significant. Statistical tests and exact sample sizes used to calculate statistical significance are stated in the figure legends. The investigators were blinded to assess all the staining assays.

### Study Approval

Human studies: The NPH_1 protocol on the research of therapeutic targets in the frame of nephronophthisis and renal associated ciliopathies has been approved by the French National Committee for the Protection of Persons (CPP) under the ID-RCB number 2016-A00541-50 and is kept in full accordance with the principles of the Declaration of Helsinki and Good Clinical Practice guidelines. The human renal biopsies belonging to the Imagine Biocollection are declared to the French Minister of Research under the number DC-2020-3994 and this biocollection was approved by the French Ethics committee for research at Assistance Publique-Hôpitaux de Paris (CERAPHP) under the IRB registration number #00011928.

Animal studies: All animals were handled in strict accordance with good animal practice as defined by the national animal welfare bodies, and all animal procedures were performed in accordance with protocols approved by the ethical committee of the « Ministère de l’Enseignement Supérieur, de la Recherche et de l’Innovation » (N°APAFIS#22608-2018042715366694 v4).

## Author contributions

H.G., A.S. and F.L. are first co-authors assigned by alphabetic order. F.L., P.C.R., S.S-M., J-P.A., A.B., M.D., L.B-R., S.S. designed research studies. H.G., F.L., A.S., E.P., C.M., A.V., E.B., M.G-T., S.R., S.C., M.M., B.D., L.F., F.J-H., M.D. conducted experiments. H.G., F.L., A.S., E.P., C.M., A.V., E.B., M.G-T., S.R., S.C., M.M., B.D., L.F., F.J-H., M.D. acquired data. H.G., F.L., A.S., E.P., C.M., A.V., E.B., M.G-T., S.R., S.C., M.M., P.C.R., B.D., L.F., S.S-M., A.B., M.D., L.B-R., S.S. analyzed data. N.C., E. B. and S.S performed RNAseq analyses. E.D.N conducted Biophenics experiments. S.S. initiated the project. H.G., K.B., and S.S. managed legal affairs. H.G., K.B., M.F., C.A., S.L., P.K., R.S. recruited subjects and/or provided clinical information and obtained all clinical samples. H.G., F.L., A.B., M.D., L.B-R. and S.S. wrote the manuscript. A.S., E.P., A.V., P.C.R., B.D., S.S-M. and J-P.A. reviewed and edited the manuscript.

## Acknowledgements

We are grateful to the patients and their families for their participation. We thank Olivia BOYER, Aurélie HUMMEL and Guillaume LEZMI (Pediatric Nephrology, Necker Enfants Malades, APHP, Paris, France), Amélie LEZMI RYCKEWAERT and Sophie TAQUE (Pediatric Hematology and Oncology, Hôpital Universitaire, Rennes, France), Odile BOESPFLUG-TANGUY (Centre de Compétence des Leucodystrophies et Leucoencéphalopathies de Cause Rare, Pôle Femme et Enfant, Hôpital Estaing, Centre Hospitalier Universitaire de Clermont-Ferrand, Clermont-Ferrand, France), Jérôme HARAMBAT/ Brigitte LLANAS (Department of Pediatrics, Bordeaux University Hospital, Bordeaux, France) who helped the follow-up of patients.

We thank the Department of Clinical Research (Salma KOTTI, Elisabeth HULIER-AMMAR) at Imagine Institute with the sponsorship team (Didier BEUDIN, Olivier FREUND, Isabelle BUFFET, Haizia HAMMOUDI, Ahlam RAMDANI, Nicholas RENAUD) that facilitates and structures the set-up of the clinical research projects and the investigation team (Pauline TOUCHE, Marc MELESAN, Aurélie JOSPIN, Joséphine CORNET, Kenza BENSARI, Maxime DOUILLET) that prepares and ensures the follow-up of the clinical trials.

We are grateful to Romain Marlange and Nicolas Doulet (Innovation & Technology Transfer Department, IMAGINE Institute) to help us to establish the collaboration with industrial partners Alexion and Medetia.

We thank Cécile Jeanpierre for critical reading of the manuscript. We would like to thank the Biophenics core facilities (Curie Institute, Paris), in particular Aurianne Lescure for her technical assistance; the zebrafish, mice, transgenesis, histology and Imaging facilities of SFR Necker, especially Pierre David for his help in establishing the *Nphp1* KO CRISPR-Cas mouse model, and Nicolas Cagnard (Bioinformatics Platform, IMAGINE Institute) for his help on RNAseq analysis; Victor Racine (QuantaCELL) for the design of the semi-quantitative analysis of zebrafish body curvature and retina layers thickness; Xavier Gérard and Jean-Michel Rozet (IMAGINE Institute) for their precious and helpful advices for phenotypic analysis of retinal degeneration. We are grateful to Michael Perricone who initiate the collaboration between IMAGINE and Alexion.

This work was supported by the Institut National de la Santé et de la Recherche Médicale (INSERM), the Ministère de l’Education Nationale de la Recherche et de la Technologie (MRT), by a State funding from the Agence Nationale de la Recherche under “Investissements d’avenir” program (ANR-10-IAHU-01) and by a public grant “RHU-C’IL-LICO” overseen by the French National Research Agency (ANR) as part of the second “Investissements d’Avenir” program (reference: ANR-17-RHUS-0002). We acknowledge the Imagine Institute for the purchase of Leica SP8 STED and Zeiss Spinning Disk microscopes, and the Fondation ARC (EML20110602384) for the purchase of the LEICA SP8 confocal microscope.

## Data availability

The data that support the findings of this study are available from the corresponding author upon reasonable request.

