## Supplementary Tables for "Prostaglandin E_1_ as therapeutic molecule for Nephronophthisis and related ciliopathies"

**Supplementary table 1:** List of the 11 compounds selected amongst the 51 hits of the *in cellulo* drug screen. The molecules were chosen based on their mode of action, toxicity and physicochemical properties.

| | Name | Class | Pertinent Effect(s) | Toxicity theshold | | | Safe Concentrations |
|---|---|---|---|---|---|---|---|
| | | | | Inhibition of proliferation | Apopoptic bodies | Permeabilized cells | |
| **A** | Alprostadil | Synthetic prostaglandin | TGF-β inhibitor, decreased cilia beat, actin & RhoA inhibitor, canonical Wnt agonist | ≥ 40µM | None up to 80µM | ≥ 80µM | < 40µM |
| **B** | Cyproheptadine hydrocloride | First-generation H1 antihistamine | CyclinD2 expression inhibitor, decreased cilia beat, apoptosis inhibitor | ≥ 5µM | ≥ 20µM | ≥ 40µM | < 5µM |
| **C** | Ethopropazine hydrocloride | Muscarinic receptor antagonist | Autophagy | ≥ 10µM | ≥ 80µM | ≥ 80µM | < 10µM |
| **D** | Fluticasone propionate | Corticosteroid | Decreased cilia beat, Smo agonist | NA* | ≥ 40µM | NA* | < 40µM |
| **E** | Methotrexate | Antimetabolite | Actin stress fibers disassembly | None up to 80µM | None up to 80µM | None up to 80µM | Up to 80µM |
| **F** | Mycophenolic Acid | Inosine monophosphate dehydrogenase inhibitor | JNK activator, MAPK/NF-κB inhibitor | None up to 80µM | ≥ 40µM | None? | < 40µM |
| **G** | Paclitaxel | Spindle poison | Microtubule depolymerization inhibitor | ≥ 10nM | ≥ 100nM | ≥ 100nM | < 10nM |
| **H** | Pyrimethamine | Antiprotozoal | STAT3 inhibitor | ≥ 10µM | None up to 80µM | ≥ 20µM | < 10µM |
| **I** | Simvastatin | HMG CoA Reductase inhibitor | RhoGTPases inhibition, actin modulation | ≥ 1µM | ≥ 1µM | ≥ 1µM | < 1µM |
| **J** | Tropisetron hydrochloride | Serotonin 5-HT3 receptor antagonist | JNK inhibitor, increased tight-junction formation, antifibrotic (skin) | ≥ 40µM | ≥ 80µM | None up to 80µM | < 40µM |
| **K** | Verapamil | Non-Dihydropyridine Calcium channel blocker | RhoGTPases inhibition & cAMP elevation, mTor, autophagy inducer | None up to 80µM | ≥ 5µM? | None up to 80µM | Up to 80µM |

* False positive beyond 5µM

**Supplementary table 2:** Genotype and clinical details of individuals from whom URECs were derived.

| | Genotype status | Clinical Status | Renal Failure (Stage) | Sex | Age (y) | % Ciliated cells |
|---|---|---|---|---|---|---|
| **Pt1** | del Hom *NPHP1* | *Nephronophthisis* | 4 | M | 26 | 19 |
| **Pt2** | del Hom *NPHP1* | *Nephronophthisis* | 4 | M | 15 | 32 |
| **Pt3** | del Hom *NPHP1* | *Nephronophthisis* | 3 | F | 15 | 21 |
| **Pt4** | Het c.1884+1G>T + Het c.1252-2A>G *NPHP1* | *Nephronophthisis* | 4 | F | 13 | 50 |
| **Pt5** | del Hom *NPHP1* | *Nephronophthisis* | 4 | M | 18 | 38 |
| **Pt6** | del Hom *NPHP1* | *Nephronophthisis* | 4 | F | 16 | 40 |
| **Pt7** | del Het + Het c.1078C>T - p.Q360* *NPHP1* | *Nephronophthisis* | 1 | M | 9 | 43 |
| **Pt8** | del Hom *NPHP1* | *Nephronophthisis* | 1 | M | 23 | 49 |
| **Pt9** | del Het + Het c.70-1G>A *NPHP1* | *Nephronophthisis* | 4 | F | 8 | 32 |
| **Pt10** | del Hom *NPHP1* | *Nephronophthisis* | 4 | F | 17 | 45 |
| **Pt11** | del Hom *NPHP1* | *Nephronophthisis* | 4 | M | 30 | 47 |
| **CTL1** | | Minimal change disease | / | F | 9 | 65 |
| **CTL2** | | No renal anomalies | / | F | 14 | 39 |
| **CTL3** | | No renal anomalies | / | F | 21 | 66 |
| **CTL4** | | No renal anomalies | / | M | 10 | 74 |
| **CTL5** | | No renal anomalies | / | M | 9 | 75 |
| **CTL6** | | No renal anomalies | / | F | 11 | 62 |
| **CKD1** | | Hemolytic uremic syndrome | 5 | M | 12 | 65 |
| **CKD2** | | Detrusor urethral sphincter dyssynergia | 4 | M | 17 | 87 |
| **CKD3** | | Twin-to-twin transfusion syndrome | 2 | F | 14 | 57 |
| **CKD4** | | Right renal hypoplasia/left renal dysplasia/bilateral vesicoureteral reflux | 4 | M | 18 | 89 |
| **CKD5** | | Renal hypodisplasia with repeated urinary infections | 3 | F | 20 | 64 |

Pt: *NPHP1* patient, CTL: control individual, CKD: non-ciliopathy chronic kidney disease patients, del: deletion, Hom: Homozygous, Het: heterozygous, M: male, F: female

**Supplementary table 3**: Genes modulated by Alprostadil treatment and NPHP1 (128 genes) (Excel file).

**Supplementary table 4:** Genes modulated by Alprostadil treatment and belonging to cilium modulators (65 genes) (Excel file).

**Supplementary table 5:** Primers used for RT-qPCR.

| Gene name | Species | Forward primer (5' to 3') | Reverse primer (5' to 3') |
|---|---|---|---|
| *Col1a1* | Mouse | CTGACGCATGGCCAAGAAGA | GGGACCCTTAGGCCATTGTG |
| *Col3a1* | Mouse | GGACCAGCAGGAACTAATGGTAT | GTTCTCCAGGTGATCCATCTTT |
| *Lcn2* | Mouse | TCCTCAGGTACAGAGCTACAA | GCTCCTTGGTTCTTCCATACA |
| *Kim1* | Mouse | GAGAGTGACAGTGGTCTGTATTG | TCATAGCAGCCACCTTCATTC |
| *Acta2* | Mouse | CATGCGTCTGGACTTGGCTG | GACAATCTCACGCTCGGCAGTAG |
| *Tgfβ1* | Mouse | GGGAAGCAGTGCCCGAACCC | TGGGGGTCAGCAGCCGGTTA |
| *Nphp1* | Mouse | GTGAGCATCAGCAGGAAAGA | CTTCTTCCTCTCCACCAGTTTC |
| *Nphp1* (genotyping) | Mouse | ACCGTTTAGTGCAACCTGTG | CATCACGTCTTCCATGCACT |
| *THBS1* | Human | CCCAGATCAGGCAGACACAG | TACTGGCAGTTGTCCCGTTC |
| *LAMC2* | Human | CGCAGCTCTGCAGAATACAG | AGACCCATTTCGTTGGACAG |
| *ITGA2* | Human | GCAACATCCCAGACATCCCA | CTTTCGTAGCACTTCGTCGC |
| *HAS3* | Human | TACATCCAGGTGTGCGACTC | CCTACTTGGGGATCCTCCTC |
| *FLNA* | Human | GTGCCAGCCGAATTCAGTAT | CACATAAGCCACACCACAGG |
| *ABLIM3* | Human | CAGTCCATGGCCAGCAGTAA | CACTTGAAGCAGCTGACGTG |
| *PKIA* | Human | TGATATCCTGGTTTCCTCTGCA | GGCTTCCCCACTTTGTTCTG |
| *CPT1A* | Human | CCATGCCATCCTGCTTTACA | ATCGTGGATCCCAAAAGACG |
| *PGC1A* | Human | CCAAAGGATGCGCTCTCGTTCA | CGGTGTCTGTAGTGGCTTGACT |
| *DEPTOR* | Human | CAGCATGTGTCCAACAAGCA | GGGGGACTTTTCATTGAGCA |
| *ATOH8* | Human | TGACTACAGTGCCGACCACA | TCACTCCTTGCGCTTCTTGG |
