## Supplementary Figures for "Prostaglandin E_1_ as therapeutic molecule for Nephronophthisis and related ciliopathies"

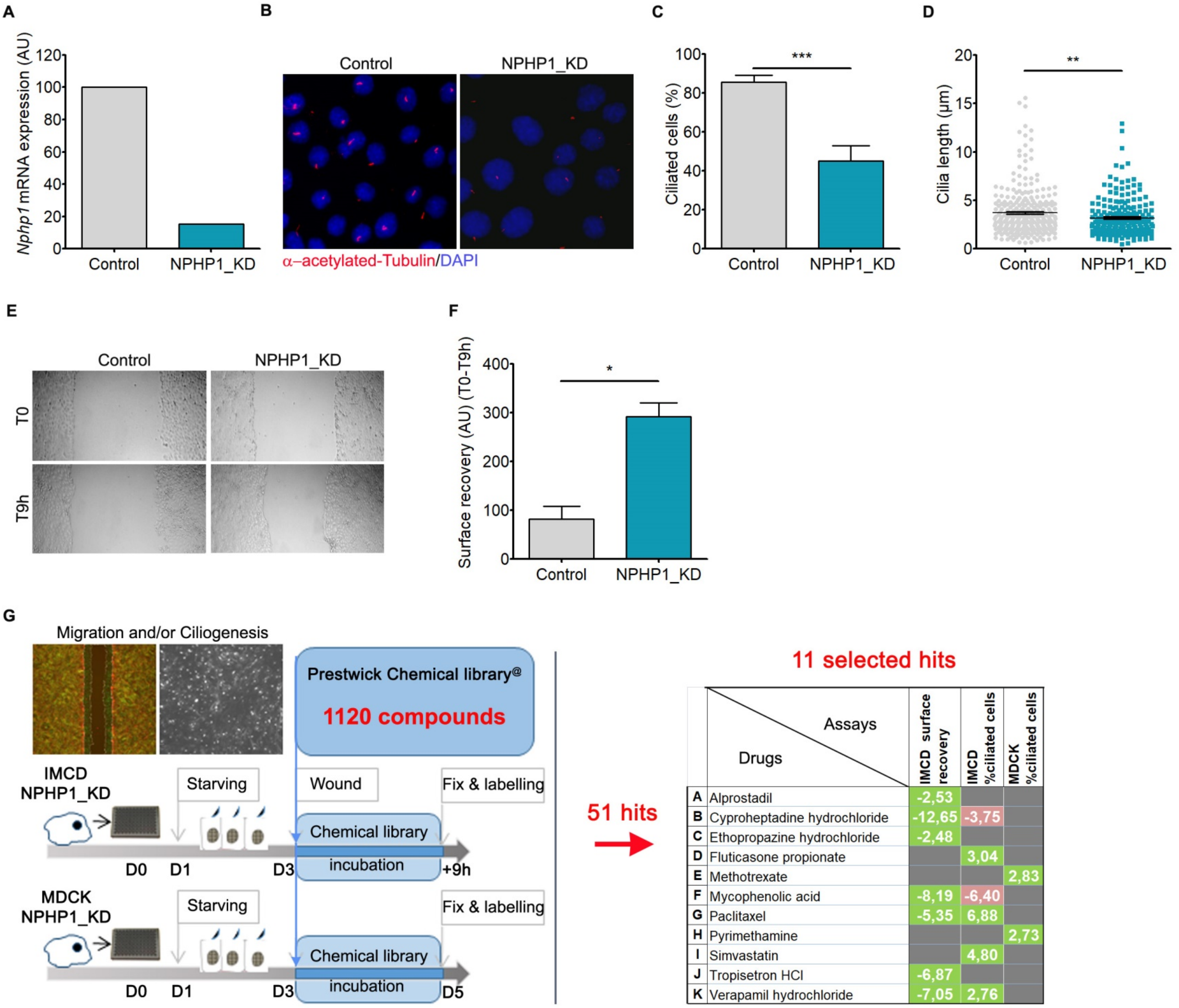


**Supplementary Figure S1: Ciliogenesis and migration defects in NPHP1_KD mIMCD-3 cells**

**(A)** RT-qPCR analysis of *Nphp1* expression in polyclonal control and *Nphp1* knock-down (NPHP1_KD) mIMCD3 cells. **(B)** Representative images of immunofluorescence of primary cilia (acetylated α-Tubulin, red) and nuclei (DAPI, blue) in control and NPHP1_KD mIMCD3 cells cultured for 48 hours after serum starvation. **(C)** Quantification of the percentage of ciliated cells. $n$ = 2 experiments. Fisher's exact test: ***$P$ < 0.001. **(D)** Quantification of cilium length. $n$ = 2 experiments. Student t-test: *$P$ < 0.01. **(E)** Representative images of wound healing assay at T = 0h and T = 9h after scratch on control and NPHP1_KD mIMCD3 cells cultured for 18 hours after serum starvation. **(F)** Quantification of scratch surface recovery as seen in (E). $n$ = 2 experiments. Mann Whitney test: *$P$ < 0.05. **(G)** Left panel: Schematic representations of the primary screens. Molecules from the Prestwick library were tested for their effects on migration and/or ciliogenesis in mIMCD3 and MDCK cells, in which the expression of *Nphp1* was stably knockdown (NPHP1_KD). In mIMCD-3 cells, compounds were added for 9h after wounding and both migration and ciliogenesis were measured. In MDCK cells, compounds were added for 2 days and ciliogenesis was analyzed. Right panel: Quantification of the effect of the 11 selected hits on cell migration (surface recovery) or ciliogenesis (% of ciliated cells). RZ score were calculated for each compound when compared to DMSO treatment. A compound is identified as a 'positive hit' (green) if its RZ-score is above 2 (for ciliogenesis) or below -2 (surface recovery) for both replicate experiments. A non-significant effect of a compound on ciliogenesis or cell migration will be indicated by a RZ-score between -2 and 2, whereas a negative ciliogenesis effect (red) will be indicated by a RZ score < -2. **(C-D, F)** Bars indicate mean ± SEM. AU: arbitrary unit.

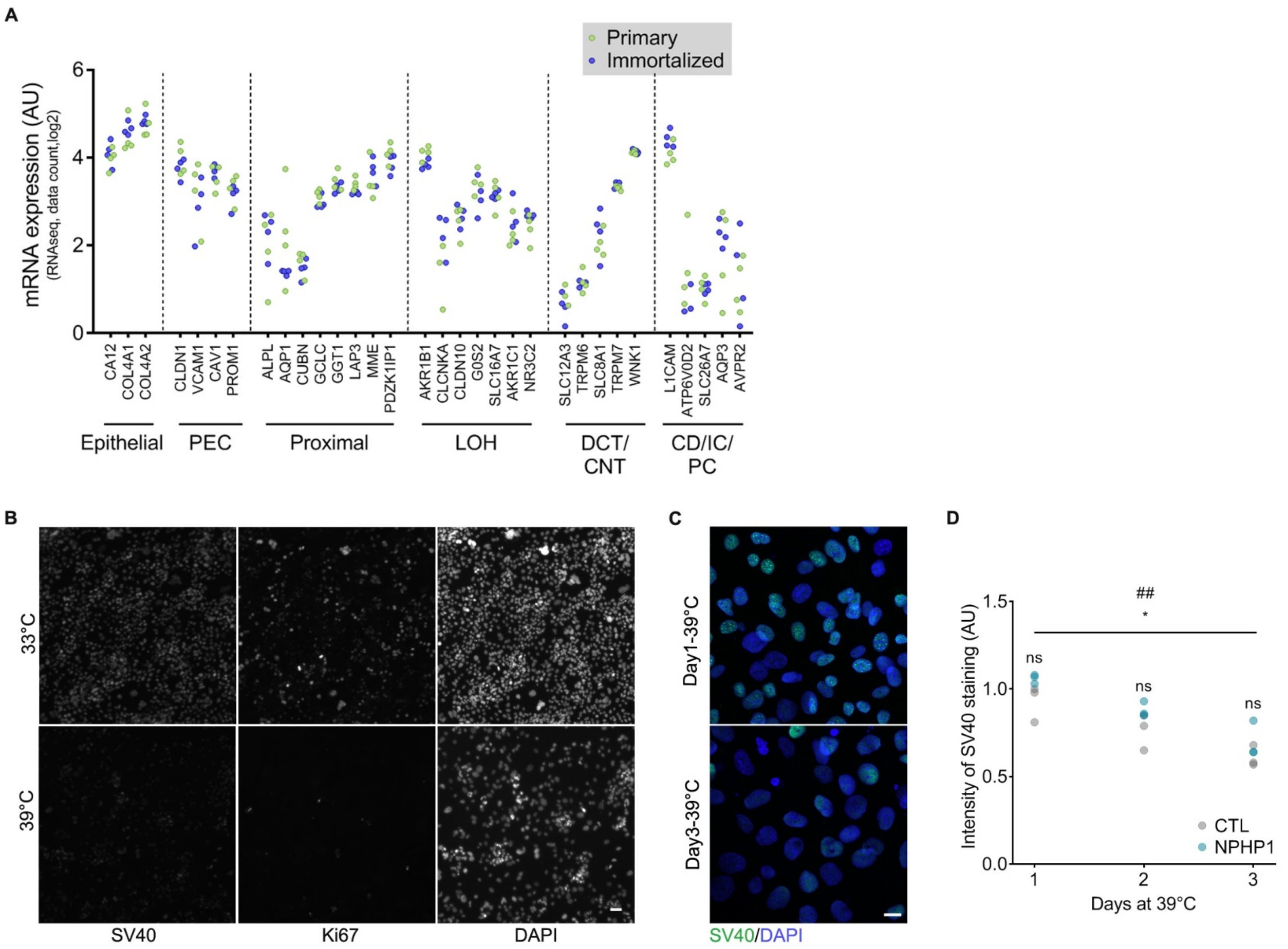


**Supplementary Figure S2: Characterization of URECs**

**(A)** Expression data of epithelial and renal markers issued from comparative RNAseq analysis of primary (green) and immortalized (blue) URECs from 4 individuals (2 NPHP1 patients and 2 control individuals). No significant *P* value observed between primary and immortalized URECs. AU: arbitrary unit. **(B)** Representative images of immunofluorescence of one immortalized UREC line derived from NPHP1 patient (Pt1) cultured for five days at 33°C (permissive temperature) or 39°C (restrictive temperature) and stained with anti-SV40 large T antigen and anti-Ki67 antibodies. **(C)** Representative images of immunofluorescence of immortalized URECs cultured for one to three days at 39°C (restrictive temperature) and stained with anti-SV40 large T antigen (green) antibody and nuclei (DAPI, blue). **(D)** Quantification of SV40 large T antigen nuclear expression in immortalized URECs derived from NPHP1 patients (NPHP1, $n = 3$) and control individuals (CTL, $n = 3$). $n = 2$ experiments. Unpaired Student t-test: *$P < 0.05$ (NPHP1 day 1 vs NPHP1 day 3), $^{\#\#}P < 0.01$ (CTL day 1 vs CTL day 3), ns: not significant (NPHP1 vs CTL). AU: arbitrary unit. Dots indicate mean. **(A, D)** Each dot represents one individual cell line. **(B, C)** Scale bar: 10µm.

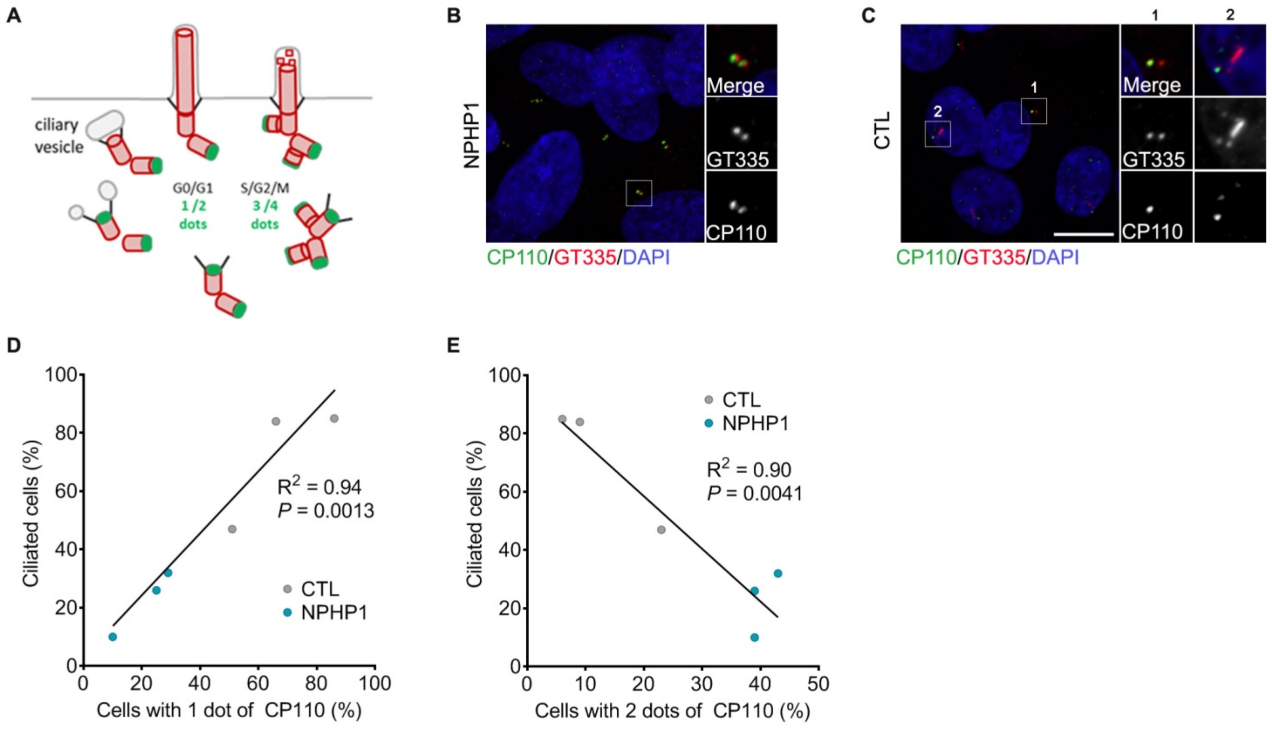


**Supplementary Figure S3: Characterization of the URECs ciliary defect**

**(A)** Scheme of CP110 cell cycle localization during ciliogenesis and cilia disassembly. **(B, C)** Representative images of immunofluorescence of immortalized URECs grown for five days at 39°C and stained for CP110 (green), cilia and basal bodies (GT335, red) and nuclei (DAPI, blue). Exemple of cells with centrosomes presenting with two (B) and one CP110 dot (C). Scale bar: 10µM. **(D, E)** Correlation between the percentage of ciliated cells and the percentage of cells with one (D) or two dots (E) of CP110. Dots indicate mean. *n* = 3 experiments. Pearson's test (*r* = 0.9700 for one dot, *r* = -0,9471 for two dots). Each dot represents one individual cell line.

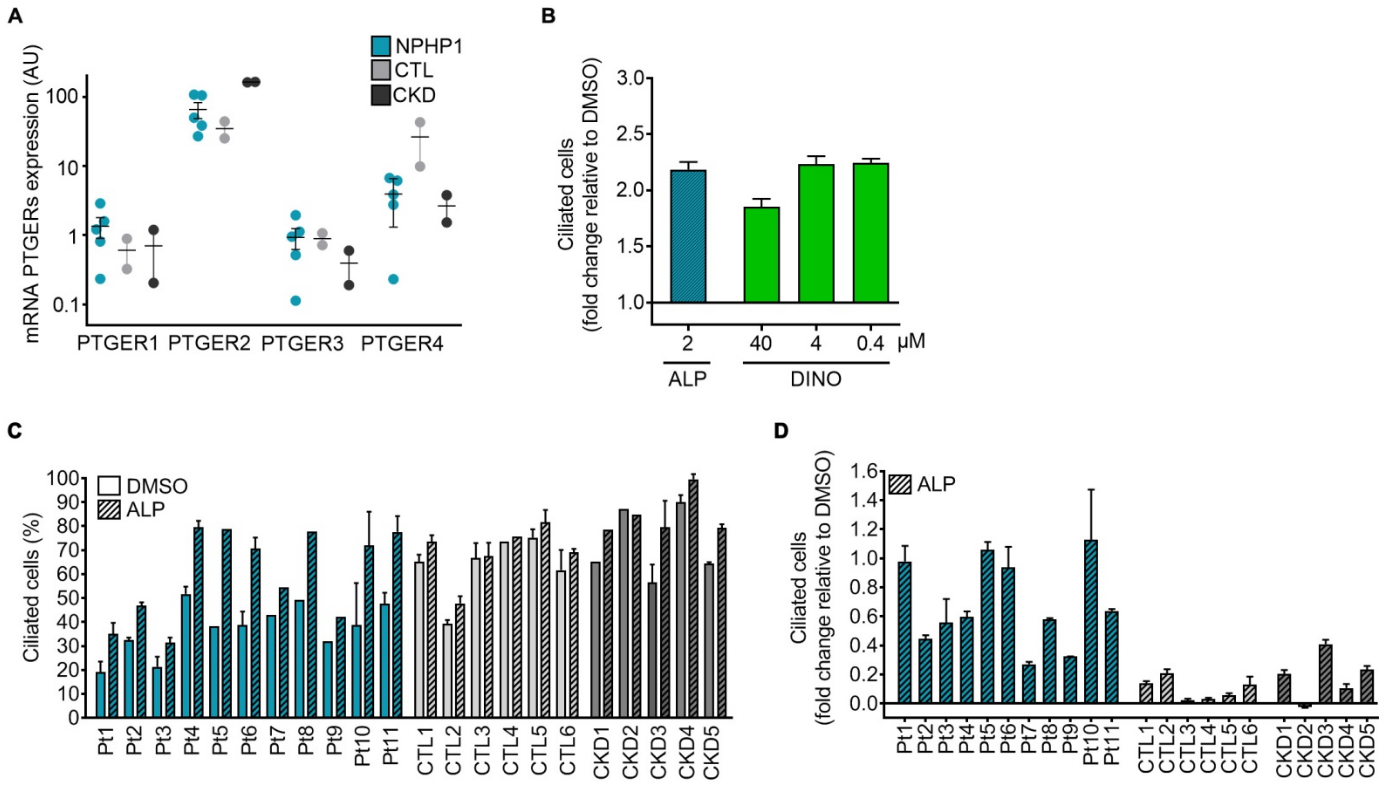


**Supplementary Figure S4: Validation of prostaglandin signaling as a target in *NPHP1* URECs**

**(A)** RT-qPCR analysis of EP receptors (*PTGER1-4*) expression in immortalized NPHP1 (NPHP1, *n* = 5), CTL (*n* = 2) and CKD (*n* = 2) URECs. AU: arbitrary unit. **(B)** Quantification of ciliogenesis 5 days after seeding in one immortalized NPHP1 UREC line (Pt1) exposed for 2 days to 2μM Alprostadil or increasing concentrations of Dinoprostone. Bars indicate mean ± SEM. *n* = 2 experiments. **(C-D)** Effect of Alprostadil on ciliogenesis, evaluated by the percentage of ciliated cells (C) or the fold change of ciliated cell relative to DMSO (D) in immortalized URECs derived from NPHP1 patients (Pt1-11), control individuals (CTL1-6) and CKD patients (CKD1-5). ALP: Alprostadil, DINO: Dinoprostone.

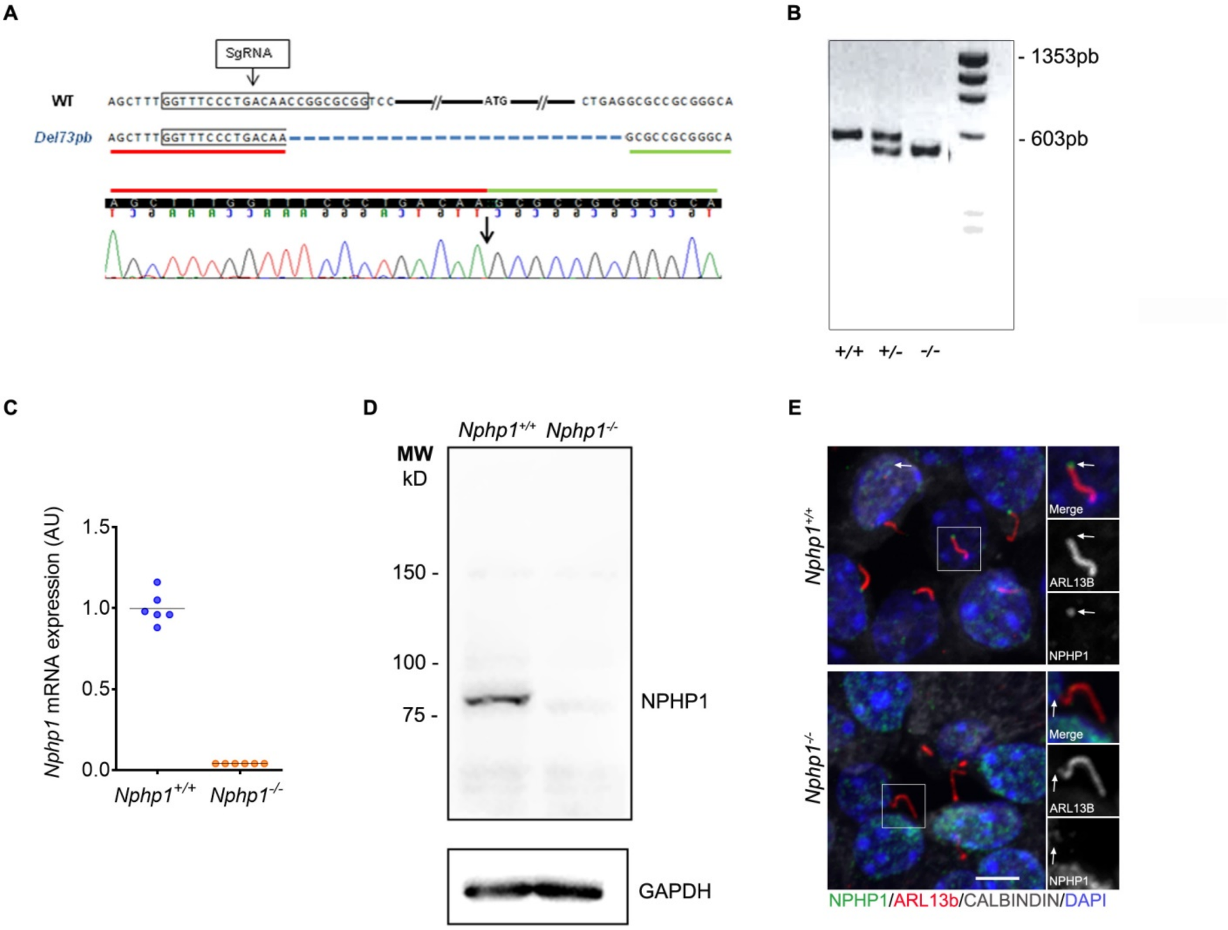


**Supplementary Figure S5: *Nphp1*$^{-/-}$ mice generation using CRISPR/Cas9 technology**

**(A)** Schematic representation of the deletion strategy to obtain *Nphp1*$^{-/-}$ mice using CRISPR/Cas9. Guide RNAs targeted sequence upstream exon 1 leads to a 73bp deletion encompassing the start codon ATG in exon 1 (Del 73pb) (Sanger Sequencing). **(B)** Genotypes were based on PCR amplification of total genomic DNA with primers framing the deletion (PCR fragment of 603pb for *Nphp1*$^{+/+}$). **(C)** RT–qPCR analysis of *Nphp1* expression in kidneys from *Nphp1*$^{+/+}$ and *Nphp1*$^{-/-}$ 5 months-old mice. Each dot represents one individual mouse. Bars indicate means. AU: arbitrary unit. **(D)** Western blot analysis of NPHP1 expression in *Nphp1*$^{+/+}$ or *Nphp1*$^{-/-}$ mice testis. **(E)** Representative images of immuostaining of NPHP1 (green), primary cilia (ARL13B, red), and nuclei (DAPI, blue) in *Nphp1*$^{+/+}$ or *Nphp1*$^{-/-}$ mice. Scale bar: 10µM.

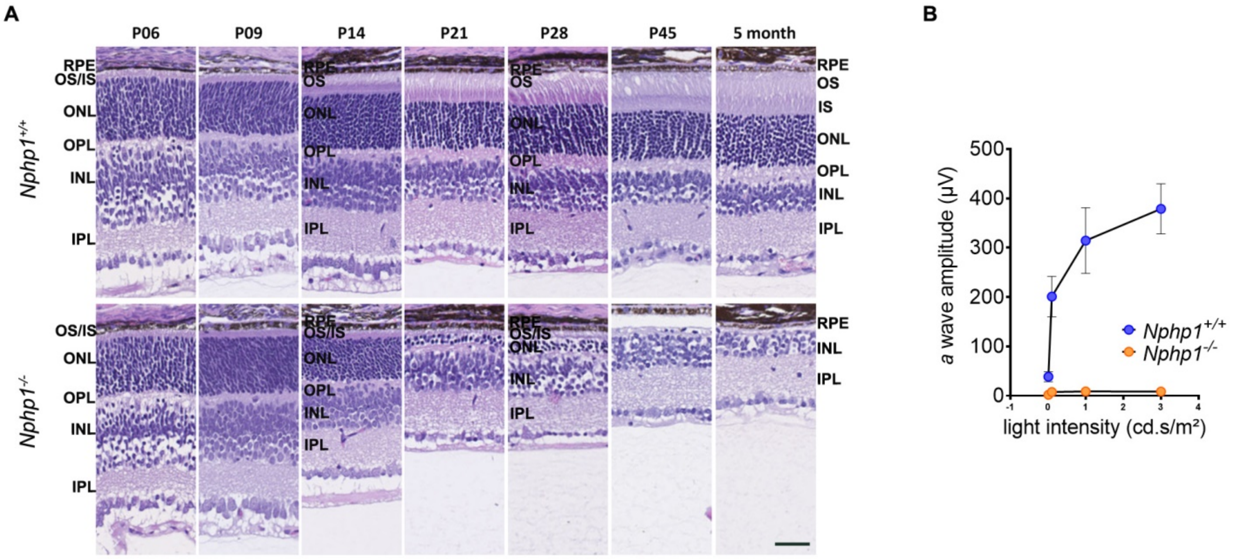


**Supplementary Figure S6: Retina phenotype of *Nphp1*$^{-/-}$ mice**

**(A)** H&E staining of *Nphp1*$^{+/+}$ and *Nphp1*$^{-/-}$ retinas from P6 to P45 and 5 months-old mice. Scale bar: 50µm. **(B)** Scotopic *a*-wave amplitude *versus* stimulus intensity (log) function in *Nphp1*$^{+/+}$ (*n* = 4) and *Nphp1*$^{-/-}$ (*n* = 4) mice retinas at P21. AU: arbitrary unit. RPE: retinal pigmented epithelium, OS outer segment, IS: inner segment, ONL: outer nuclear segment, OPL: outer plexiform layer, INL: inner nuclear segment, IPL: inner plexiform layer.

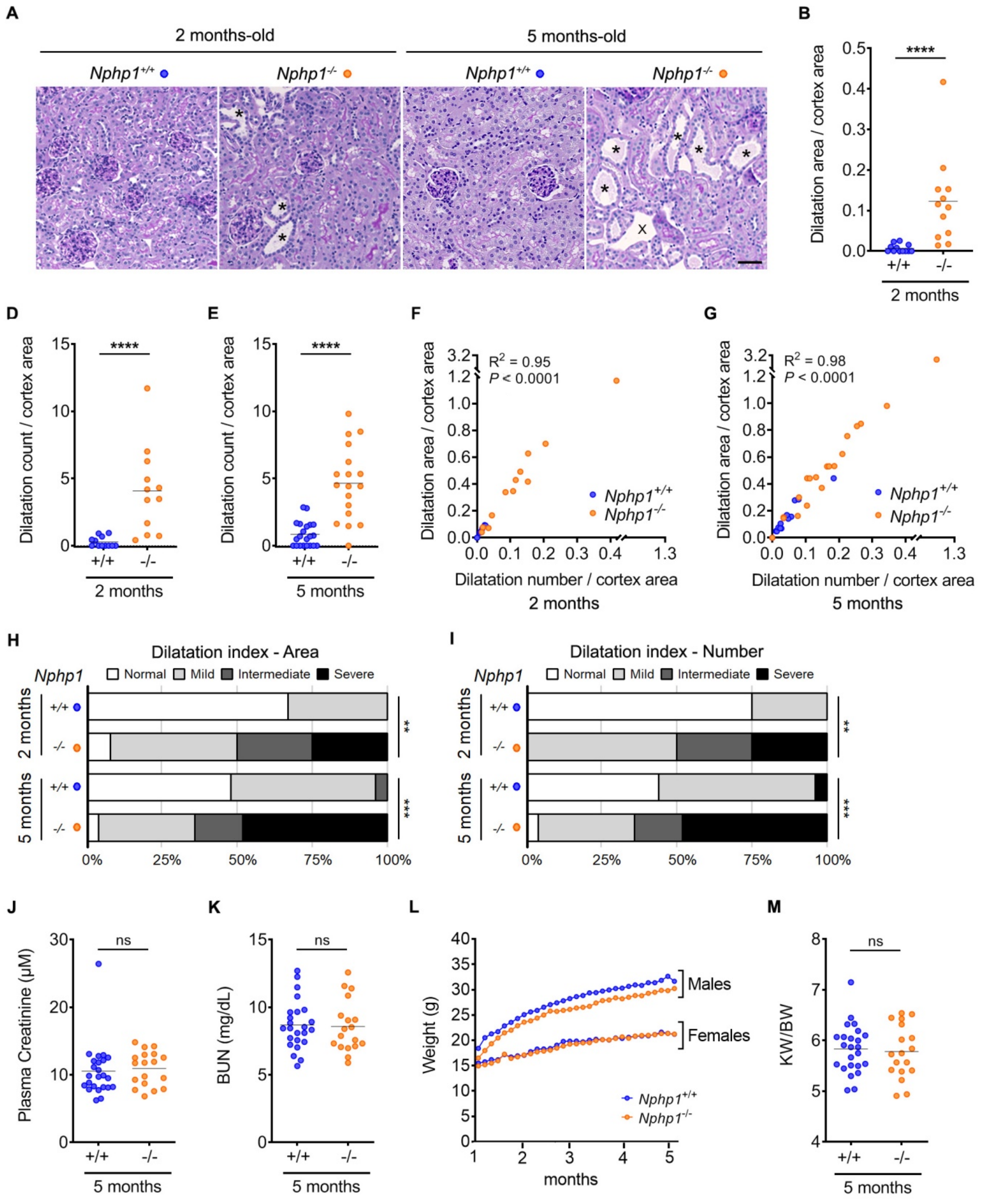


**Supplementary Figure S7: Analysis of kidney phenotype from *Nphp1*$^{+/+}$ and *Nphp1*$^{-/-}$ mice**

**(A)** Representative images of PAS staining of kidney sections from 2 and 5 months-old *Nphp1*$^{+/+}$ and *Nphp1*$^{-/-}$ mice. Scale bar: 50µm. * = tubular dilatations; X = blood vessels. **(B-E)** Quantification of the total area covered by dilatation/cortex area (B, C) and the number of dilatation/10µm$^2$ in cortex (D, E) in *Nphp1*$^{+/+}$ and *Nphp1*$^{-/-}$ mice at 2 and 5 months-old. Mann Whitney test: ****$P < 0.0001$. **(F, G)** Scatter dot plots correlating the number of dilatation/10µm$^2$ in cortex to the total area covered by dilatations/cortex area in 2 months (F) and 5 months (G) -old *Nphp1*$^{+/+}$ mice. Pearson's test ($r$ = 0,9770 for 2 months-old, $r$ = 0.9914 for 5 months-old). **(H, I)** Dilatation index for the total area covered by dilatations/cortex area (H) and the number of dilatation/10µm$^2$ in cortex (I). Chi-squared test: **$P < 0.01$, ***$P < 0.001$. **(J, K)** Plasma creatinine (J) and blood urea nitrogen (BUN) (K) of 5 months-old *Nphp1*$^{+/+}$ and *Nphp1*$^{-/-}$ mice. Mann Whitney test: ns: not significant. **(L)** Body weight increase in *Nphp1*$^{+/+}$ and *Nphp1*$^{-/-}$ mice from 1 month to 5 months-old. **(M)** Kidney weight to body weight ratio (KW/BW) of 5 months-old *Nphp1*$^{+/+}$ and *Nphp1*$^{-/-}$ mice. Mann Whitney test: ns: not significant. **(B-M)** 2 months-old: males ($n$ = 6), females ($n$ = 6), 5 months-old: males ($n$ = 12)), females ($n$ = 12)) and *Nphp1*$^{-/-}$ (2 months-old: males ($n$ = 7), females ($n$ = 5), 5 months-old: males ($n$ = 8), females ($n$ = 10). **(B-G, J, K, M)** Each dot represents one individual mouse.

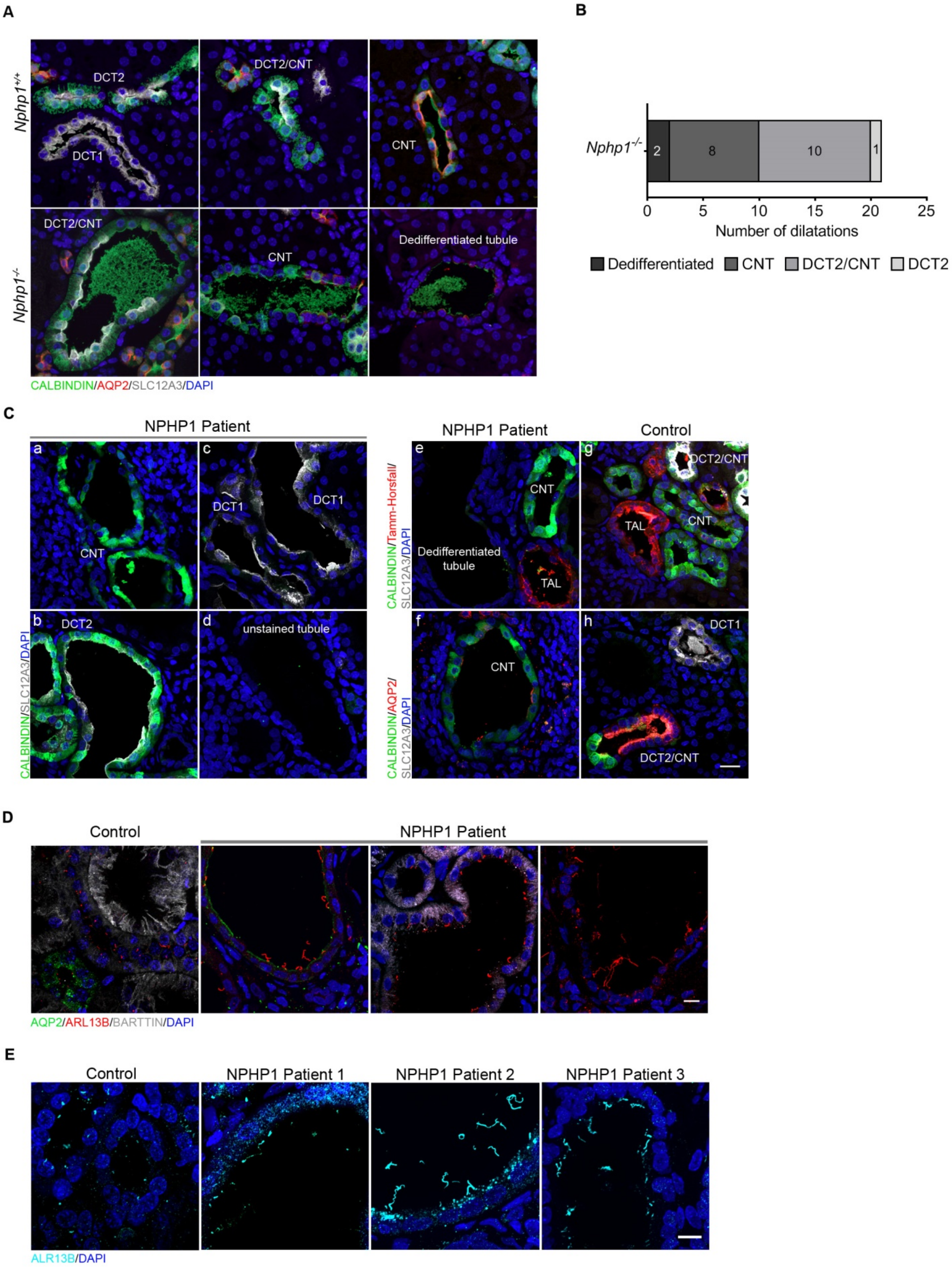


**Supplementary Figure S8: Immunohistological analyses of *Nphp1*$^{-/-}$ mouse and *NPHP1* patient kidneys**

**(A)** Representative images of immunostaining of mouse kidney tubules stained with anti-SLC12A3 (DCT, gray), anti-Calbindin (DCT and CNT, green), anti-AQP2 (CNT and CCD, red) antibodies and DAPI (nuclei, blue). Scale bar: 100µm. **(B)** Quantification of the number of dilatations in each segment of the nephron in *Nphp1*$^{-/-}$ animals ($n$ = 4). **(C)** Representative images of (a-d) immunostaining of human kidney tubules from *NPHP1* patient (NPHP1) stained with anti-SLC12A3 (gray), anti-Calbindin (green) antibodies and DAPI (blue); (e-g) immunostaining of human kidney tubules from *NPHP1* patient (e) and control individual (g) with anti-SLC12A3 (gray), anti-Calbindin (green), anti-Tamm-Horsfall (TAL, red) antibodies and DAPI (blue); (f-h) immunostaining of human kidney tubules from *NPHP1* patient (f) and control individual (h) with anti-SLC12A3 (gray), anti-Calbindin (green), anti-AQP2 (red) antibodies and DAPI (blue). Scale bar: 20µm. **(D)** Representative images of immunostaining of human kidney sections from NPHP1 patient and control individual stained with anti-AQP2 (green), anti-ARL13B (cilia, red), anti-BARTTIN (TAL, DCT and CNT, gray) antibodies and DAPI (blue). Scale bar: 10µm. **(E)** Immunostaining of a control and three additional NPHP1 patient's kidney sections stained with anti-ARL13B (cyan) antibody and DAPI (nuclei, blue). Scale bar: 20µm. TAL: Thick Ascending Limb of the Loop of Henle, DCT: Distal Convoluted Tubule, CNT: Connecting Tubule, CCD: Cortical Collecting Duct.

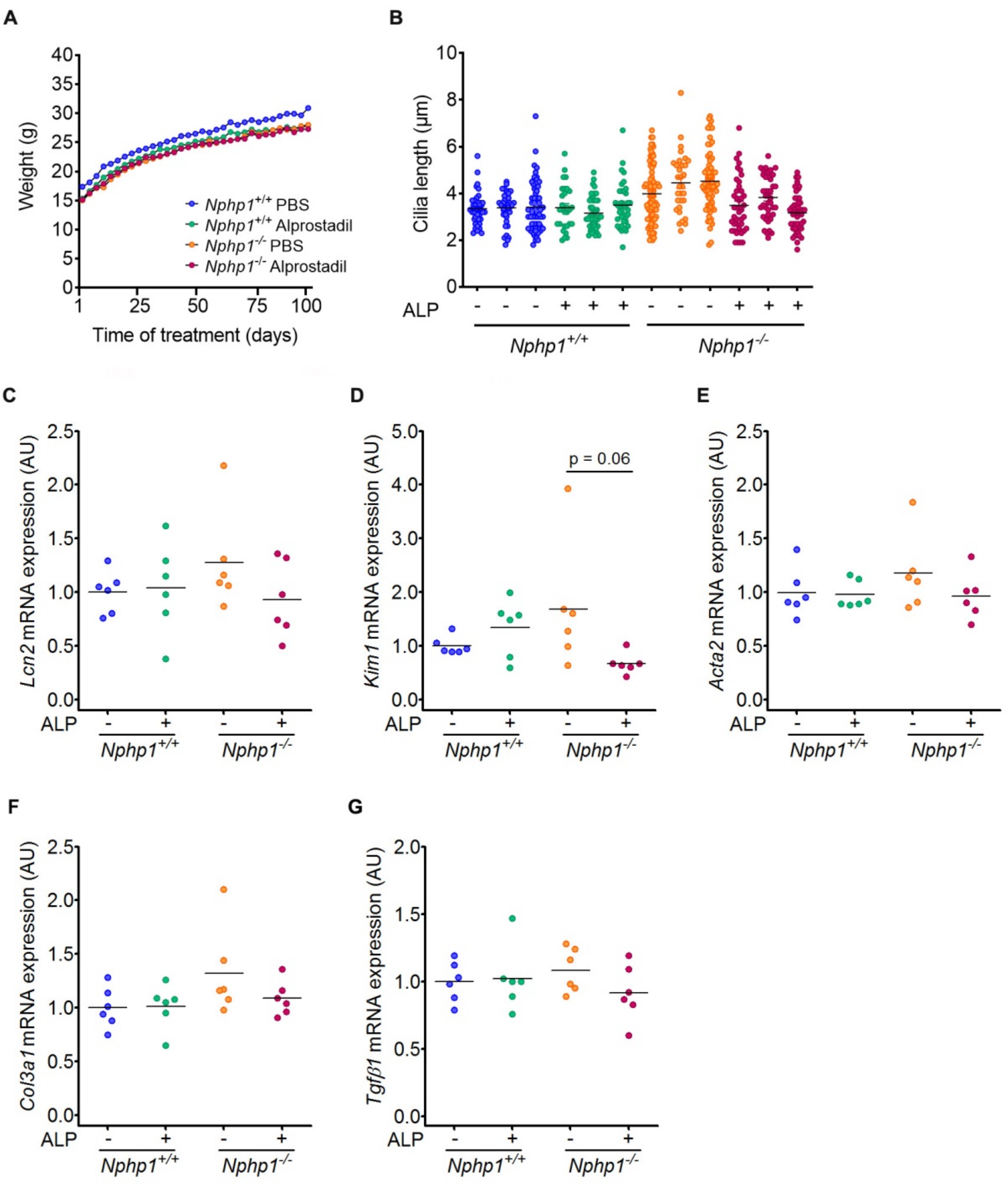


**Supplementary Figure S9: Impact of Alprostadil treatment on ciliogenesis and pro-fibrotic markers in kidney of *Nphp1*$^{+/+}$ and *Nphp1*$^{-/-}$**

**(A)** Body weight increase in *Nphp1*$^{+/+}$ and *Nphp1*$^{-/-}$ mice injected daily with vehicle (PBS) or Alprostadil (80µg/kg) from 1 month to 5 months-old (*n* = 9-10 male mice for each genotype/treatment). **(B)** Effect of Alprostadil (ALP) on primary cilium length in connecting and distal convoluted tubules of *Nphp1*$^{+/+}$ and *Nphp1*$^{-/-}$ mice at 5 months-old. Each dot represents one cilium. **(C-G)** RT-qPCR analysis of *Lcn2*, *Kim1*, *Acta2*, *Col3a1* and *Tgfβ1* expression from 5 months-old *Nphp1*$^{+/+}$ and *Nphp1*$^{-/-}$ mice kidneys treated or not with Alprostadil. One-way ANOVA followed by Holm-Sidak's post-test: not significant. Each dot represents one individual mouse. AU: arbitrary unit. **(B-G)** Bars indicate mean. ALP: Alprostadil.

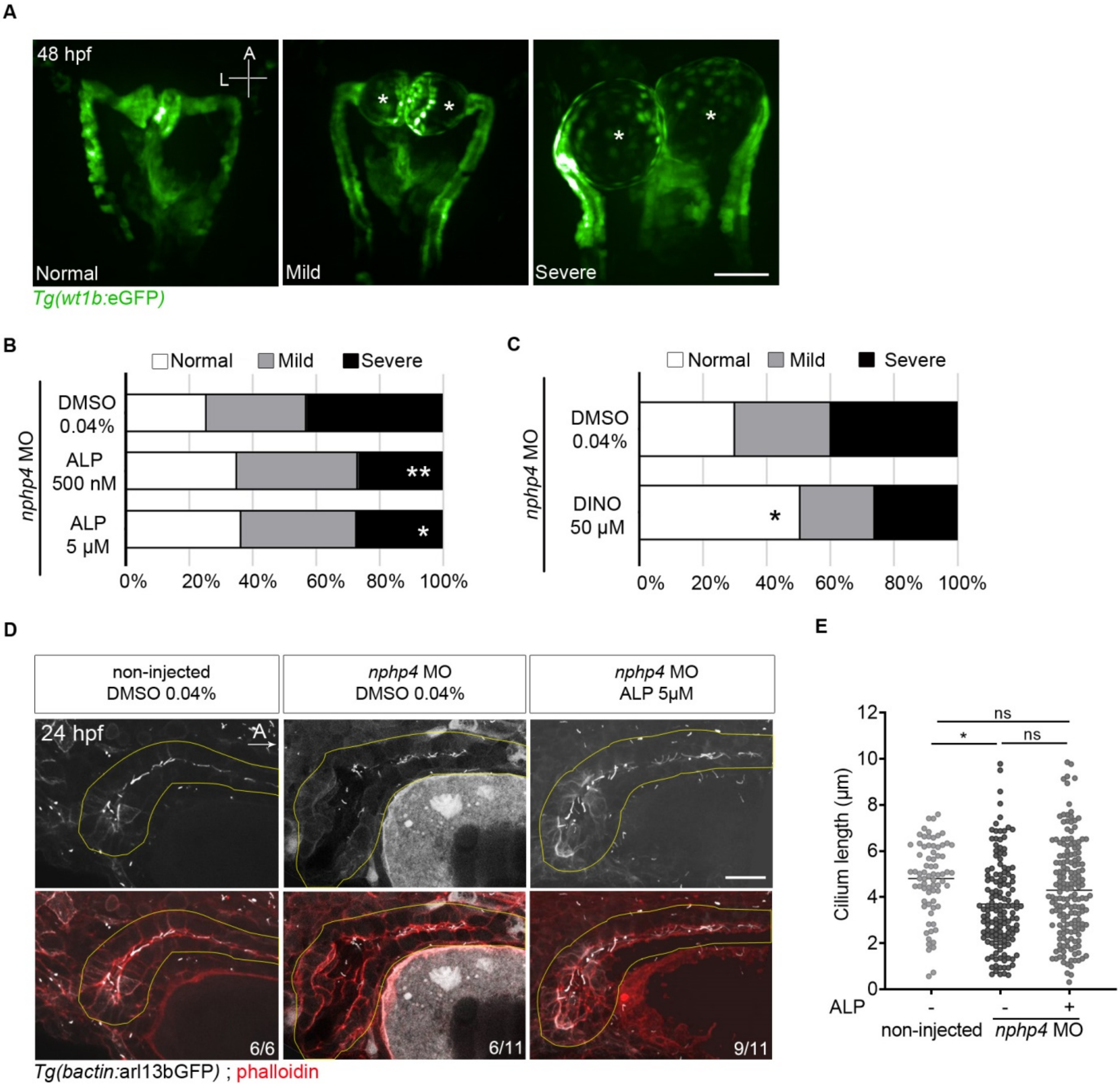


**Supplementary Figure S10: *In vivo* validation of prostaglandin signaling as a target in zebrafish**

**(A)** *nphp4* morpholino-injected embryos were treated from 10 to 48 hours post fertilization (hpf), and morphology of glomeruli and proximal pronephros was assessed using *Tg(wt1b:*GFP*)* transgenic embryos at 48-54 hpf. Three different categories were established depending of the absence (normal) or presence of mild or severe glomerular dilations (asterisks). Representative images of dorsal views, anterior to the top. Scale bar: 50µm. **(B)** Treatment with Alprostadil significantly reduced the percentage of severe dilations at 500nM (*n* = 71 embryos) and 5µM (*n* = 99 embryos) compared to control DMSO (0.04%) (*n* = 86 embryos). **(C)** Treatment with Dinoprostone (50µM) (*n* = 77 embryos) significantly increased the percentage of normal proximal pronephros compared to DMSO (0.04%) (*n* = 93 embryos). **(D)** *nphp4*-morpholino injected embryos were treated with Alprostadil (5µM) from 10 to 24 hpf and fixed prior to immunostaining. Tubule lumens were labelled with phalloidin, and cilia with *Tg(bactin:arl13bGFP)* transgene. *nphp4* knockdown leads to dilation of the cloaca region, that is partially restored with Alprostadil treatment. Representative images of side views, anterior to the right. Scale bar: 20µm. **(E)** Quantification of cilium length in the distal part of the pronephros of *nphp4* morpholino-injected embryos upon Alprostadil (*n* = 14 embryos) or DMSO (*n* = 11 embryos) treatment, compared to DMSO-treated non-injected animals (*n* = 6 embryos). Bars indicate mean. Mann Whitney test: **P* < 0.05, ns = not significant. Each dot represents cilia length. ALP: Alprostadil, DINO: Dinoprostone. **(B-C)** *n* = 3–5 experiments. Fisher's exact test (Normal/Mild compared to Severe (B), Normal compared Mild/Severe (C)): **P* < 0.05, ***P* < 0.01.
